# CAP2 is a novel regulator of Cofilin in synaptic plasticity and Alzheimer’s disease

**DOI:** 10.1101/789552

**Authors:** Silvia Pelucchi, Lina Vandermeulen, Lara Pizzamiglio, Bahar Aksan, Jing Yan, Anja Konietzny, Elisa Bonomi, Barbara Borroni, Marco Rust, Daniele Di Marino, Marina Mikhaylova, Daniela Mauceri, Flavia Antonucci, Fabrizio Gardoni, Monica Di Luca, Elena Marcello

**Author notes:** Co-first authors (Silvia Pelucchi, Lina Vandermeulen). Co-senior authors (Monica Di Luca, Elena Marcello). Corresponding author: Prof. Elena Marcello, Department of Pharmacological and Biomolecular Sciences, Università degli Studi di Milano, via Balzaretti 9, 20133 Milan, Italy.

## Abstract

Cofilin is one of the major regulators of actin dynamics in spines where it is required for structural synaptic plasticity. However, our knowledge of the mechanisms controlling Cofilin activity in spines remains still fragmented. Here, we describe the cyclase-associated protein 2 (CAP2) as a novel master regulator of Cofilin localization in spines. The formation of CAP2 dimers through its Cys^32^ is important for CAP2 binding to Cofilin and for normal spine actin turnover. The Cys^32^-dependent CAP2 homodimerization and association to Cofilin are triggered by long-term potentiation (LTP) and are required for LTP-induced Cofilin translocation into spines, spine remodeling and the potentiation of synaptic transmission. This mechanism is specifically affected in the hippocampus, but not in the superior frontal gyrus, of both Alzheimer’s Disease (AD) patients and APP/PS1 mice, where CAP2 is down-regulated and CAP2 dimer synaptic levels are reduced. In AD hippocampi, Cofilin preferentially associates with CAP2 monomer and is aberrantly localized in spines. Taken together, these results provide novel insights into structural plasticity mechanisms that are defective in AD.

## Introduction

Alzheimer’s Disease (AD) is a definitive neurodegenerative disorder characterized by β-amyloid (Aβ) plaques and neurofibrillary tangles. Beside this knowledge, much evidence supports synaptic dysfunction as a preceding and contributing insult to eventual neuronal death (Selkoe, 2002). Dendritic spine loss is seen in post mortem brains from AD patients and in mice models of AD (Marcello *et al*, 2012a). In AD patients, cognitive decline has a stronger correlation to synapse loss than to neurofibrillary tangles or neuronal loss (DeKosky & Scheff, 1990). Diverse lines of evidence suggest that Aβ oligomers play a prominent role in AD synaptic dysfunction since they lead to dendritic spines loss and aberrant plasticity phenomena (Lacor *et al*, 2007; Shankar *et al*, 2007; Walsh *et al*, 2002).

Dendritic spines are small dendritic protrusions containing excitatory postsynaptic machinery (Bourne & Harris, 2008). Since changes in spine morphology account for functional differences at the synaptic level (Yuste & Bonhoeffer, 2001), spine remodeling and modifications in spine density are believed to be the basis of learning and memory (Holtmaat & Svoboda, 2009) and are thereby associated with brain diseases characterized by cognitive decline (Penzes *et al*, 2011). Indeed, it is widely accepted that spines constitute the anatomical locus of plasticity, where short-term alterations in synaptic strength are converted into long-lasting changes that are embedded in stable structural modifications (Sala & Segal, 2014). Hence, spine structural plasticity is tightly coordinated with synaptic function and plasticity: spine enlargement parallels modification of the number, types and properties of surface glutamate α-amino-3-hydroxy-5-methyl-4-isoxazolepropionic acid receptors (AMPAR) during long-term potentiation (LTP), whereas during long-term depression (LTD) the decrease in synaptic strength is associated with spine shrinkage (Kasai *et al*, 2010).

In this frame, the actin cytoskeleton is important for postsynaptic structure, function and plasticity because it confers spine plasticity and stability (Cingolani & Goda, 2008; Hotulainen & Hoogenraad, 2010). Actin is highly enriched in spines and filamentous actin (F-actin) is the major structural backbone of spines since it forms organized bundles in spine necks (Star *et al*, 2002). Only a relatively small fraction of actin in spines is stable (Kasai *et al*, 2010), while the most abundant dynamic fraction of the actin cytoskeleton provides the driving force behind structural remodeling of spines and contributes to synaptic plasticity (Matus, 2005).

Regulation of actin dynamics is also relevant in AD pathogenesis, since the signaling pathways influencing actin cytoskeleton remodeling have been shown to be impaired in AD (Penzes & Vanleeuwen, 2011). In particular, among the actin-binding proteins implicated in AD pathology, Cofilin plays a critical role. Indeed, abnormalities of Cofilin have been reported in AD patients (Bamburg & Bernstein, 2016) and Aβ oligomers affect Cofilin activation (Henriques *et al*, 2015). Cofilin is a key bidirectional regulator of spine structural plasticity, as it is implicated in both spine enlargement (Bosch *et al*, 2014) and spine shrinkage (Zhou *et al*, 2004; Pontrello *et al*, 2012; Hotulainen *et al*, 2009; Rust *et al*, 2010). Cofilin controls F-actin assembly and disassembly in a complex, concentration-dependent manner (Hild *et al*, 2014; Rust, 2015b). At low concentrations, Cofilin promotes F-actin disassembly by accelerating the dissociation of monomeric actin (G-actin) from the filaments’ minus ends and by severing F-actin (Blanchoin & Pollard, 1999). Conversely, at high concentrations, Cofilin can promote F-actin assembly by nucleating new and by stabilizing preexisting filaments (Andrianantoandro & Pollard, 2006). Indeed, during LTP Cofilin is massively transported to the spine where it promotes the F-actin assembly that is required for spine expansion (Bosch *et al*, 2014).

The actin dynamizing activity of Cofilin is enhanced by binding partners as the cyclase-associated proteins (CAPs) (Normoyle & Brieher, 2012). CAPs are evolutionary highly conserved multi-domain actin binding proteins capable of regulating actin dynamics at multiple levels (Ono, 2013). CAP deficiency results in defects in vesicle trafficking, endocytosis, and in an altered cell morphology and cell growth (Noegel *et al*, 1999). Two closely related homologs of CAP have been described in mammals. CAP1 is expressed in nearly all cells, whereas CAP2 expression is restricted to a limited number of tissues, including the brain (Bertling *et al*, 2004; Peche *et al*, 2007), suggesting that CAP2 may have unique roles, particularly in neuronal cells. CAP2 gene deletion has been described in a rare developmental disorder, named 6p22 syndrome, which is characterized by developmental delays and autism spectrum disorders (Field *et al*, 2015). In addition, alterations in spine morphology and dendrite architecture have been reported in CAP2 knock-out neurons (Kumar *et al*, 2016).

Here, we introduce a novel mechanism, altered in both AD patients and APP/PS1 mice hippocampi, through which the dimerization of CAP2, dependent on Cys^32^, is relevant for actin turnover in spines and is necessary to target Cofilin to spines upon LTP induction. The mutation of CAP2 Cys^32^ is sufficient to prevent the LTP-triggered changes in spine morphology and function. These findings may open new ways in our understanding and targeting synaptic dysfunction and spine dysmorphogenesis in AD.

## Results

### Cofilin synaptic localization and association with CAP2 are impaired in Alzheimer’s disease

Although an impaired activation of Cofilin has been reported in the hippocampus of AD patients (Barone *et al*, 2014; Minamide *et al*, 2000), nothing is known about its synaptic localization in AD. Given the importance of Cofilin translocation in the spine for synaptic plasticity (Bosch *et al*, 2014; Mikhaylova *et al*, 2018), we first assessed Cofilin synaptic levels in the hippocampus of APP/PS1 mice at 6 months of age, when they begin to develop Aβ deposits (Jankowsky *et al*, 2004). We purified the Triton-insoluble fraction (TIF), which is highly enriched in postsynaptic proteins (Gardoni *et al*, 2001; Marcello *et al*, 2012b). Western Blot analysis of the total homogenate and of the TIF samples shows a significant increase of Cofilin synaptic levels in APP/PS1 mice hippocampi compared to wild-type mice, while no changes in the total protein levels were detected (Fig. 1A).

**Figure 1.**
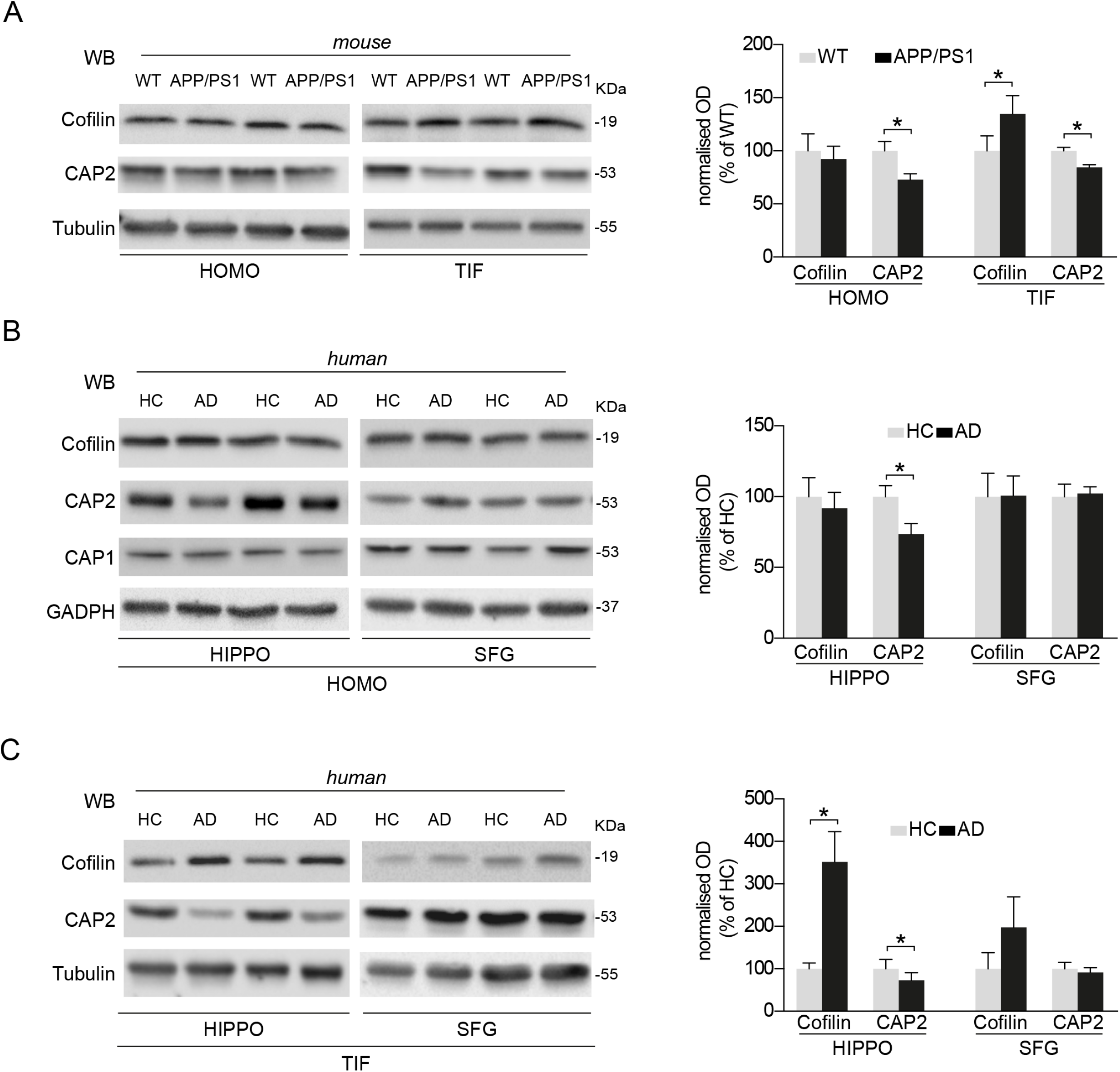
The synaptic localization of Cofilin and CAP2 is specifically altered in the hippocampus of AD patients APP/PS1 mice. A. Representative Western Blot (WB) showing the expression of Cofilin, CAP2 and Tubulin in homogenate (HOMO) and triton-insolible fraction (TIF) samples of APP/PS1 mice at 6 months of age compared to wild type (WT) animals hippocampi; quantification of optical density (OD) and normalization on loading control OD (Tubulin) shows that CAP2 but not Cofilin total levels are reduced in APP/PS1 mice (CAP2, WT vs APP/PS1 \**P* = 0.023, 2-tailed unpaired t-test, *n*=6). In the TIF fraction, the quantitative analysis reveals that Cofilin levels increase in transgenic mice, while CAP2 levels decrease (Cofilin, WT vs APP/PS1 \**P* = 0.015; CAP2, WT vs APP/PS1 \**P* = 0.026, 2-tailed paired t-test, *n*=6). B Representative Western Blot (WB) showing the expression of Cofilin, CAP2, CAP1 and GAPDH in homogenate samples (HOMO) of AD and HC hippocampi (HIPPO) and SFG; quantification of optical density (OD) and normalization on loading control OD (GAPDH) shows that CAP2 but not Cofilin total levels are reduced in AD patients (CAP2, HC vs AD \**P* = 0.027, 2-tailed unpaired t-test, *n*=9). In SFG HOMO samples Cofilin and CAP2 levels were not affected in AD patients compared to HC Cofilin, HC vs AD \**P*>0.05; CAP2, HC vs AD *P*>0.05, 2-tailed paired t-test, *n*=6). No changes in CAP1 levels in the HOMO of both HIPPO and SFG were observed (CAP1, HIPPO, HC=0.522±0.045, AD= 0.468±0.050, HC vs AD *P* >0.05, 2-tailed unpaired t-test, *n*=9; SFG, HC= 1.448± 0.077, AD= 1.604± 0.179, HC vs AD *P* >0.05, 2-tailed unpaired t-test, *n*=6) C Representative WB showing Cofilin, CAP2 and Tubulin levels postsynaptic TIF samples of the HIPPO and SFG of AD and HC. The quantitative analysis reveals that Cofilin levels increase in the TIF of AD patients while CAP2 levels decrease (Cofilin, HC vs AD \**P* = 0.02; CAP2, HC vs AD \**P* = 0.02, 2-tailed paired t-test, *n*=9). No significant modifications of Cofilin and CAP2 synaptic localization were detected in the SFG (Cofilin, HC vs AD \**P*>0.05; CAP2, HC vs AD *P*>0.05, 2-tailed paired t-test, *n*=6). In this and all subsequent figures, data represent mean ± SEM.

The protein CAP2 has been described as a novel Cofilin binding partner (Kumar *et al*, 2016). In mice, we observed that CAP2 is a protein expressed in the cortex, striatum, cerebellum and brain stem from postnatal day 0 (P0), while in the hippocampus CAP2 levels are detectable from P14 (EV1A). Interestingly, the CAP2 protein levels in the total homogenate and, consistently, in the postsynaptic fraction of APP/PS1 mice were significantly reduced compared to wild-type mice (Fig. 1A).

These data indicate an altered synaptic availability of Cofilin and of its binding partner CAP2 in the hippocampal synapses of APP/PS1 mice at the early stages of the pathology.

To strengthen these results, we took advantage of autoptic hippocampus and superior frontal gyrus (SFG) specimens obtained from sporadic AD patients, fulfilling criteria for Braak 4 and 5 stage, and age-matched control subjects (HC) (Table 1, 2). The hippocampus can be considered the first brain area affected by the disease, while SFG is a less affected area that we used as negative control because no significant plaque deposition was detectable at this stage, as revealed by neuropathology reports.

**Table1.**
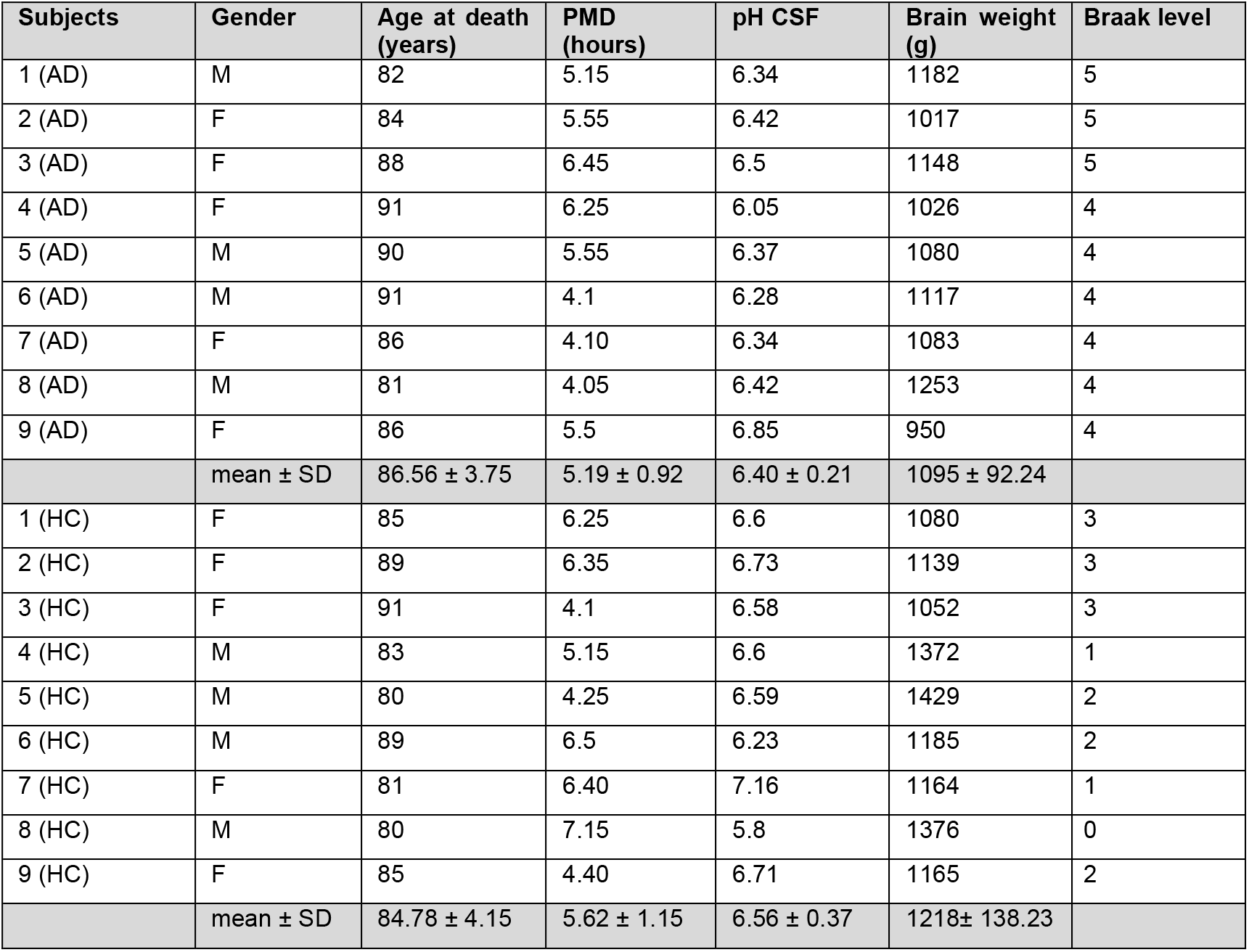
Demographic and neuropathological characteristics of AD and HC cases that were selected for the analysis of hippocampal specimens (AD, Alzheimer’s disease; CSF, cerebrospinal fluid; F, female; HC, healthy controls; M, male; PMD, postmortem delay)

**Table2.** Demographic and neuropathological characteristics of AD and HC cases that were selected for the analysis of SFG specimens (AD, Alzheimer’s disease; CSF, cerebrospinal fluid; F, female; HC, healthy controls; M, male; PMD, postmortem delay)

| Subjects | Gender | Age at death (years) | PMD (hours) | pH CSF | Brain weight (g) | Braak level |
| --- | --- | --- | --- | --- | --- | --- |
| 1 (AD) | F | 91 | 3.45 | 6.36 | 1011 | 4 |
| 2 (AD) | F | 91 | 4.15 | 6.27 | 1202 | 4 |
| 3 (AD) | F | 86 | 4.10 | 6.34 | 1083 | 4 |
| 4 (AD) | F | 86 | 5.05 | 6.62 | 998 | 4 |
| 5 (AD) | M | 81 | 4.05 | 6.42 | 1253 | 4 |
| 6 (AD) | F | 86 | 5.5 | 6.85 | 950 | 4 |
| | mean ± SD | 86.83 ± 3.76 | 4.38 ± 0.75 | 6.48 ± 0.22 | 1082 ± 120 | |
| 1 (HC) | F | 84 | 4.45 | 6.26 | 1179 | 1 |
| 2 (HC) | F | 81 | 6.40 | 7.16 | 1164 | 1 |
| 3 (HC) | M | 80 | 7.15 | 5.8 | 1376 | 0 |
| 4 (HC) | M | 84 | 7.05 | 5.9 | 1385 | 1 |
| 5 (HC) | F | 85 | 5.00 | 6.72 | 1257 | 1 |
| 6 (HC) | F | 85 | 4.40 | 6.71 | 1165 | 2 |
| | mean ± SD | 83.17 ± 2.14 | 5.74 ± 1.28 | 6.43 ± 0.53 | 1254 ± 103 | |

The analysis of the total homogenate revealed no changes in Cofilin total protein levels and a significant down-regulation of CAP2 in AD patients hippocampi (Fig. 1B), as previously reported in microarray studies (Blalock *et al*, 2004). This AD-associated alteration of CAP2 is specific since we detected no modifications in the total levels of the homolog CAP1 in AD patients’ hippocampi and SFG when compared to HC (Fig. 1B). After the validation of the postsynaptic fraction purification protocol from human specimens (EV1B), we performed Western blot analysis of the TIF samples. We measured a significant increase in Cofilin synaptic levels and a concomitant decrease in CAP2 synaptic localization (Fig. 1C), as observed in APP/PS1 hippocampi (Fig. 1A). Interestingly, these alterations are specific for the hippocampus because no modifications of CAP2 and Cofilin total levels and synaptic localization were found in SFG (Fig. 1B, C).

### CAP2 is a synaptic protein relevant for synaptic function and neuronal structure

Since the AD-related alterations of CAP2/Cofilin complex have been detected in the postsynapse, we decided to further investigate CAP2 role in neuronal cells.

First, we assessed CAP2 subcellular localization. We employed stimulated emission depletion (STED) nanoscopy to analyze primary hippocampal neurons stained for endogenous CAP2, pre- and postsynaptic markers and F-actin, labeled by phalloidin. Endogenous CAP2 has a distribution pattern similar to phalloidin, is localized throughout the dendrites and is detectable in dendritic spines, as previously reported (Kumar *et al*, 2016) (EV2A). A linescan through the spine length shows that peaks of fluorescence for CAP2 and the presynaptic marker Bassoon do not overlap, while CAP2 fluorescence profile shows a partial overlap with the postsynaptic protein PSD-95 profile (Fig. 2A) (Smith *et al*, 2014).

**Figure 2.**
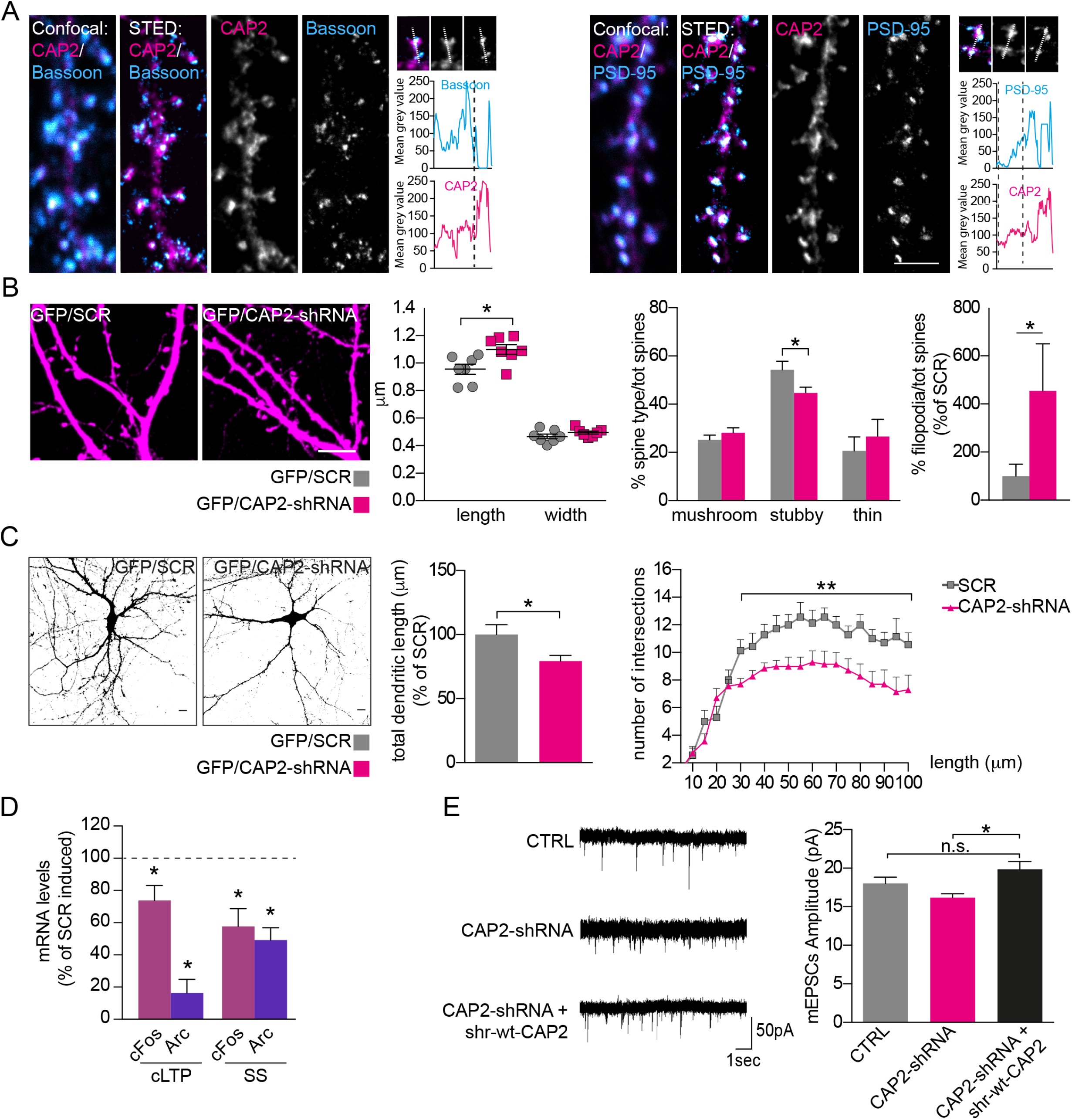
CAP2 is a postsynaptic protein critical for neuronal structure and function. A Confocal and STED images of hippocampal neurons fixed and stained for CAP2 (*magenta*) and synaptic markers (PSD-95 or Bassoon, *cyan*). Linescan through spine head was used to evaluate the mean grey value of the stained proteins. Scale bar = 2 µm. B Representative confocal images of primary hippocampal neurons transfected with GFP and either CAP2-shRNA or scramble sequence (SCR); scale bar = 5 µm. Histograms show the quantification of dendritic spine length (SCR vs CAP2-shRNA \**P* = 0.017), dendritic spine width, dendritic spine type percentage (stubby, SCR vs CAP2-shRNA \**P* = 0.035) and filopodia (SCR vs CAP2-shRNA \**P* = 0.037); 2-tailed unpaired t-test, n=7-8 neurons from at least 2 independent cultures. C Representative micrographs of hippocampal neurons transfected with GFP and either CAP2-shRNA or SCR. Scale bar = 10 µm. Total dendritic length analysis (SCR vs CAP2-shRNA \**P* = 0.04, 2-tailed unpaired t-test) and Sholl analysis (**, *P*<0.01, one-way ANOVA, uncorrected Fisher’s LSD); *n*=7-8 neurons from at least 2 independent cultures. D QRT-PCR analysis of cFos and Arc expression in hippocampal neurons infected with either rAAV-CAP2-shRNA or rAAV-SCR and with or without treatment to induce either cLTP or synaptic stimulation (SS). The treated samples are normalized on either the cLTP-SCR or the SS-SCR condition. The down-regulation of CAP2 prevents the induction of Arc and cFos upon cLTP and SS (cLTP-SCR vs cLTP CAP2-shRNA: cFos \**P*=0.048, Arc \**P*=0.013; SS-SCR vs SS-CAP2-shRNA: cFos \**P*=0.018, Arc \**P*=0.003, 2-tailed paired t-test, *n*=5 independent experiments). E (*Left*) Representative traces of excitatory miniature postsynaptic currents (mEPSCs) recorded from 15 DIV hippocampal neurons obtained from CTRL, CAP2-shRNA and CAP2-shRNA+shr-wt-CAP2 transfected cultures. (*Right*) Electrophysiological analysis of mEPSCs amplitude (pA): \**P*=0.013, one-Way ANOVA followed by Tukey’s multiple comparison test.

To further confirm the localization of CAP2 in the postsynaptic compartment, we performed biochemical fractionation of rat hippocampi to purify the postsynaptic densities (PSD). Western Blot analysis demonstrated that CAP2 is present in the synaptic fractions but not enriched in highly detergent-insoluble fractions PSD2, which corresponds to the ‘core’ of the PSD (EV2B).

To gain insights into the specific role of CAP2 in hippocampal neuronal cells and considering the reduction of CAP2 protein levels in the hippocampi of AD patients and APP/PS1 mice (Fig. 1A, B), we took advantage of small hairpin RNA (shRNA) to down-regulate CAP2 expression. We first identified a shRNA that reduced by about 80% the levels of co-transfected Myc-CAP2 in heterologous cells (CAP2-shRNA, EV2C). The same shRNA reduced by about 80% endogenous CAP2 mRNA levels when packaged into adeno-associated virus (rAAV) particles and used to infect primary hippocampal neurons (EV2D).

Since CAP2 is a postsynaptic actin binding protein, we investigated whether its down-regulation affects spine morphology and synaptic transmission in hippocampal neurons.

The analysis of the spine shape of hippocampal neurons transfected with CAP2-shRNA and GFP revealed a significant increase in spine length, while no changes in head width or density of spines were detected (Fig. 2B, SCR: 2.61±0.30 spines/10µm; CAP2-shRNA: 2.65±0.36 spines/10µm). This effect on spine length goes along with a decrease in the stubby spines population and an increase of filopodia (Fig. 2B), leading to altered neuronal spine morphology. Notably, in primary hippocampal neurons transfected with CAP2-shRNA and GFP, to visualize dendrite architecture, the Sholl analysis revealed a reduction in the complexity of the dendritic tree. (Fig. 2C). This effect was accompanied by a significant reduction in total dendritic length (Fig. 2C). Interestingly, the down-regulation of CAP2 levels also reduces the mRNA levels of BDNF, a neurotrophin that controls dendrite morphology (Horch & Katz, 2002) (EV2D). These data indicate that the down-regulation of CAP2 affects neuronal architecture and spine morphology. Therefore, we asked whether the decreased CAP2 expression could influence synaptic function. Firstly, we observed that the knockdown of CAP2 impairs the expected increase in the transcription of immediate early genes (cFos and Arc) after the induction of both chemical LTP (cLTP) and synaptic stimulation (SS) (Fig. 2D). This indicates the relevance of CAP2 in synaptic plasticity phenomena and is consistent with an alteration of neuronal architecture leading to impaired integration of signal-regulated transcription. Secondly, we recorded the miniature excitatory postsynaptic currents (mEPSCs) in neurons that revealed impairments in the post-synaptic AMPAR sensitivity as indicated by the lower mEPSCs amplitude in CAP2-shRNA neurons respect to control conditions (Fig. 2E). To assess whether this effect could be specifically ascribed to CAP2, we generated and validated a shRNA-resistant (shr) ‘rescue’ form of Myc-CAP2 (shr-wt-CAP2) that was insensitive to the CAP2-shRNA (EV2E). As shown in Fig 2E, the mEPSCs amplitude was restored to control levels by the shr-wt-CAP2, demonstrating the specificity of the effect on synaptic transmission. These results indicate that the down-regulation of CAP2 leads to a less responsive/functional postsynaptic compartment.

Collectively these data suggest that CAP2 is a postsynaptic protein relevant for shaping dendritic and spine morphology as well as for the regulation of synaptic transmission and plasticity, which are all deeply linked to the actin polymerization through Cofilin activity (Hotulainen *et al*, 2009; Gu *et al*, 2010).

### CAP2 forms disulfide-crosslinked dimers

The importance of protein self-association has been previously demonstrated for Srv2/CAP, the yeast homologous form of CAP2 (Ono, 2013; Hubberstey & Mottillo, 2002). Indeed, Srv2/CAP complex catalytically accelerates Cofilin-dependent actin turnover (Balcer *et al*, 2003). Therefore, we wondered whether the mammalian CAP2 is also able to dimerize and oligomerize. Different experimental approaches were used to test this hypothesis. Either Myc-CAP2 or EGFP-CAP2 were transfected in COS-7 cells and we observed that both constructs are able to form aggregates in the cytoplasm, independently of the tag (Fig. 3A, lower panels). The transfection of both CAP2 constructs in COS-7 cells demonstrates that Myc-CAP2 and EGFP-CAP2 perfectly colocalize in cytoplasmic clusters, suggesting that CAP2 can self-associate (Fig. 3A). Coimmunoprecipitation assays performed from lysates of COS-7 cells cotransfected with Myc-CAP2 and EGFP-CAP2 confirmed that Myc-CAP2 interacts with EGFP-CAP2 (Fig. 3B).

**Figure 3.**
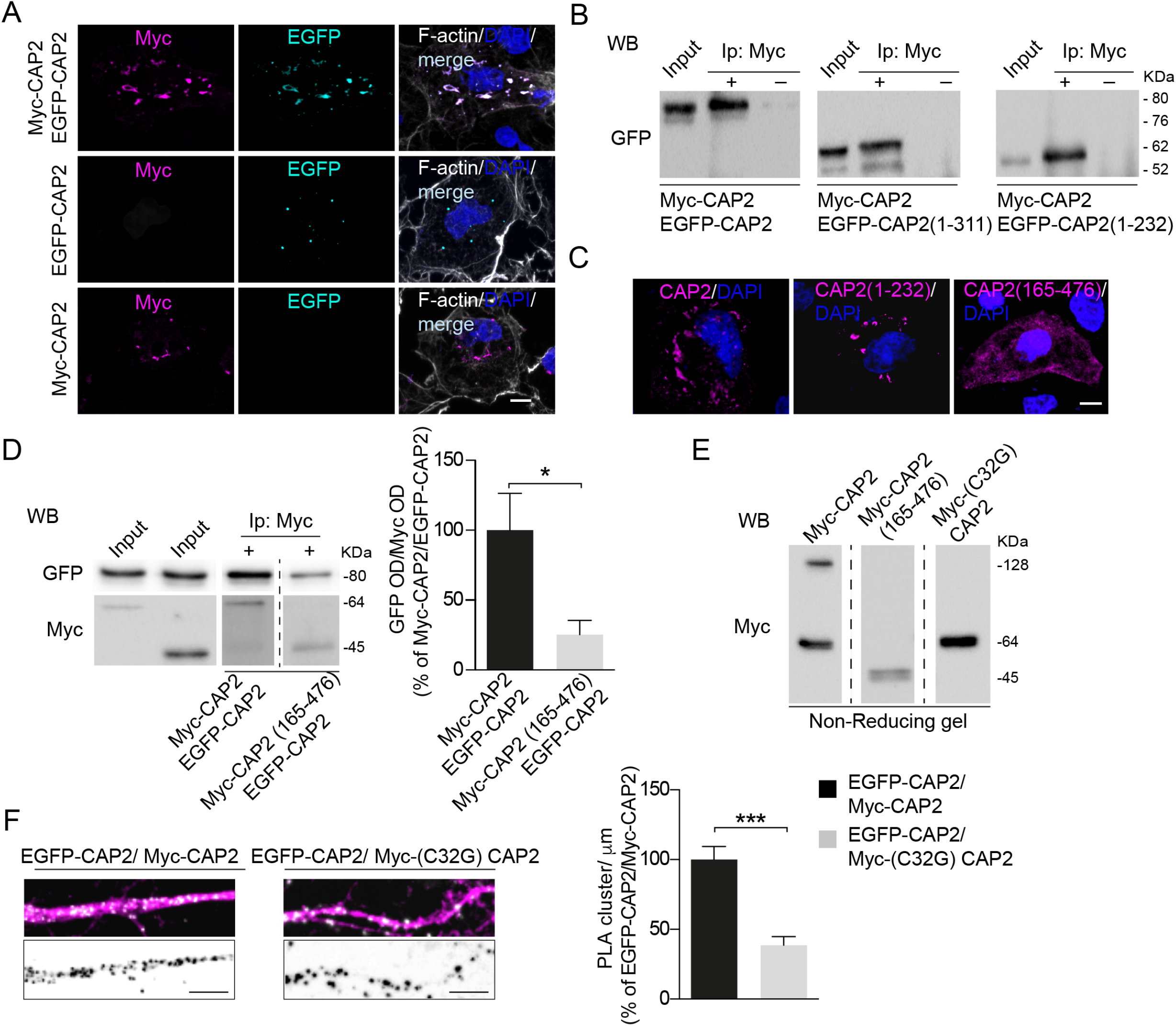
CAP2 self-associates and forms dimers through Cys^32^. A Representative confocal images of COS7 cells co-transfected with either Myc-CAP2 (*magenta*) or EGFP-CAP2 (*cyan*) or both constructs show the capability of CAP2 to form clusters. Scale bar = 10 µm. B Homogenate of HEK293 cells transfected with Myc-CAP2 and either EGFP-CAP2 or EGFP-CAP2(1-232) or EGFP-CAP2(1-311) deletion mutants were immunoprecipitated (Ip) with anti-Myc antibody and the Western Blot analysis (WB) carried out with anti-GFP antibody. The sequence 1-232 of CAP2 is sufficient for CAP2 self-association. C Representative confocal images of COS7 cells transfected with Myc-CAP2, Myc-CAP2(1-232) and Myc-CAP2(165-476) reveal that the region 1-232 is sufficient for clusters formation while the deletion of the region 1-164 completely changes the protein intracellular pattern. Scale bar = 5 µm. D Co-immunoprecipitation assays carried out from homogenate of HEK293 cells transfected with EGFP-CAP2 and either Myc-CAP2 or Myc-CAP2(165-476). The analysis of the optical density (OD) reveals that the lack of the sequence 1-164 of CAP2 significantly reduces CAP2 self-association (\**P*=0.031, 2-tailed unpaired t-test, *n* = 3-4 independent experiments, the samples derive from the same experiment and gels/blots were processed in parallel). E Representative WB of homogenates of HEK293 cells expressing Myc-CAP2, Myc-CAP2(165-476) or Myc-(C32G)CAP2. The deletion of the 1-164 region or the single point mutation Cys^32^ to Gly avoids the capability to form dimers. The samples derive from the same experiment and gels/blots were processed in parallel. F Representative confocal images of PLA experiments showing proximity between EGFP-CAP2 and either Myc-CAP2 or Myc-(C32G)CAP2 (*white*) along MAP2 positive dendrites (*magenta*); lower panels, inverted images of PLA signal (*black*), scale bar = 5 µm (\*\*\**P*=0.0002, 2-tailed unpaired t-test, *n* = 16 neurons from 2 independent experiments).

The full three-dimensional (3D) structure of the human CAP2 is not available, but several studies have underlined the multi-domains organization of the protein (Ono, 2013). The only structural information that can be useful to propose a possible 3D organization of the CAP2 dimer comes from the X-ray structure of the Helical Folded Domain (HFD) of the CAP homolog from *Dictyostelium discoideum* (Yusof *et al*, 2005). In details, this region is composed of an initial Coiled-coil region followed by a stable bundle of six antiparallel α-helices (HFD domain) (EV3A). This region (i.e. Coiled-coil and HFD) is not only conserved in CAP1 and CAP2 from human and mouse, but also in the homologous CAP protein from *Dictyostelium discoideum* (EV3B). This evidence strengthens the idea that the dimerization/oligomerization of CAP2 is crucial for its cellular functions, such as the interaction with other protein partners (i.e. actin and Cofilin).

To better characterize the region of CAP2 for the self-association, we transfected COS-7 cells with Myc-CAP2 and either EGFP-CAP2 or the deletion mutant EGFP-CAP2(1-311) deleted of the CARP domain or the EGFP-CAP2(1-232) mutant lacking of the CARP domain, the proline rich sequences and the WH2 region (Peche *et al*, 2013) (EV3A). Coimmunoprecipitation experiments revealed that CAP2 self-association depends on its N-terminal region, similarly to what previously described for Srv2/CAP (Quintero-Monzon *et al*, 2009) (Fig. 3B). To investigate the involvement of N-terminus in CAP2-self-association, we performed imaging analysis in COS-7. The deletion mutant lacking the coiled-coil region and HFD, Myc-CAP2(165-476), displayed a diffuse staining throughout the cell, while Myc-CAP2 and Myc-CAP2(1-232) showed a clustered cytoplasmic distribution (Fig. 3C). In addition, coimmunoprecipitation assays confirmed that the deletion of the region 1-164 significantly reduces the self-association (Fig. 3D).

It has been recently demonstrated that CAP1 homodimerization depends on the formation of a disulfide bond that requires the Cys^29^ of CAP1 (Liu *et al*, 2018). Therefore, we verified the presence of a disulfide cross-linked CAP2 dimer by loading a lysate of COS-7 cells transfected with Myc-CAP2 onto a non-reducing SDS-PAGE. As shown in Fig. 3E, both a band corresponding to Myc-CAP2 monomer at 64 kDa and a species of CAP2 with an apparent molecular weight of approximately double the Myc-CAP2 monomer (128 kDa) were detected. The deletion of the N-terminal region [Myc-CAP2(165-476)] abolishes the CAP2 capability to form dimers (Fig. 3E), demonstrating that this is the domain involved in the dimerization. Interestingly, only one Cys residue is present in the CAP2 region spanning from amino acid 1 to 164. Notably, the Myc-CAP2 mutant carrying the mutation of the Cys^32^ to Gly [Myc-(C32G)CAP2] is not able to dimerize in non-reducing SDS-PAGE (Fig. 3E). To strengthen these data, we performed a proximity ligation assay (PLA) in hippocampal neurons transfected with EGFP-CAP2 and either Myc-CAP2 or Myc-(C32G)CAP2. In neurons expressing EGFP-CAP2 and Myc-CAP2, a large number of PLA signals were detected when the two antibodies anti-GFP and anti-Myc were used, indicating that these two CAP2 constructs are in close proximity to each other along MAP2-positive dendrites (Fig. 3F). In neurons transfected with EGFP-CAP2 and the mutant Myc-(C32G)CAP2, the density of PLA signals significantly decreased when compared to cells overexpressing EGFP-CAP2 and Myc-CAP2 (Fig. 3F). For control experiments, only the anti-Myc primary antibody was used and no PLA signal was generated (EV3C). Overall, these findings indicate that the Cys^32^ is fundamental for the formation of CAP2 dimers and for CAP2 clustering in neurons.

### CAP2 dimer binds Cofilin and is essential for actin turnover in spines

To investigate the direct effect of monomeric and dimeric CAP2 on the assembly and stability of individual actin filaments, we employed *in vitro* dual-color total internal reflection fluorescence (TIRF) microscopy. *In vitro* polymerization of a mixture of unconjugated and Alexa-Fluor-561-conjugated actin occurs spontaneously at certain actin concentrations until it reaches equilibrium of G- and F-actin. We then added the purified recombinant CAP2 and (C32G)CAP2 proteins and determined the rate of F-actin depolymerization caused by G-actin washout, using kymograph analysis. The data showed a clear increase of actin depolymerization in the presence of CAP2 that was prevented by the mutation of Cys^32^ to glycine (Fig. 4A). In addition, the analysis of the filaments showed that the CAP2-dependent increase in actin depolymerization rate was related to an increase in the severing activity (Fig. 4B). Non-reducing SDS-PAGE assays of the recombinant proteins confirmed that CAP2, but not the mutant (C32G)CAP2, forms dimers (EV4A). Remarkably, Western blot analysis revealed that Cofilin was co-purified with CAP2, but in a smaller extent with the mutant (C32G)CAP2 (Fig. 4C).

**Figure 4.**
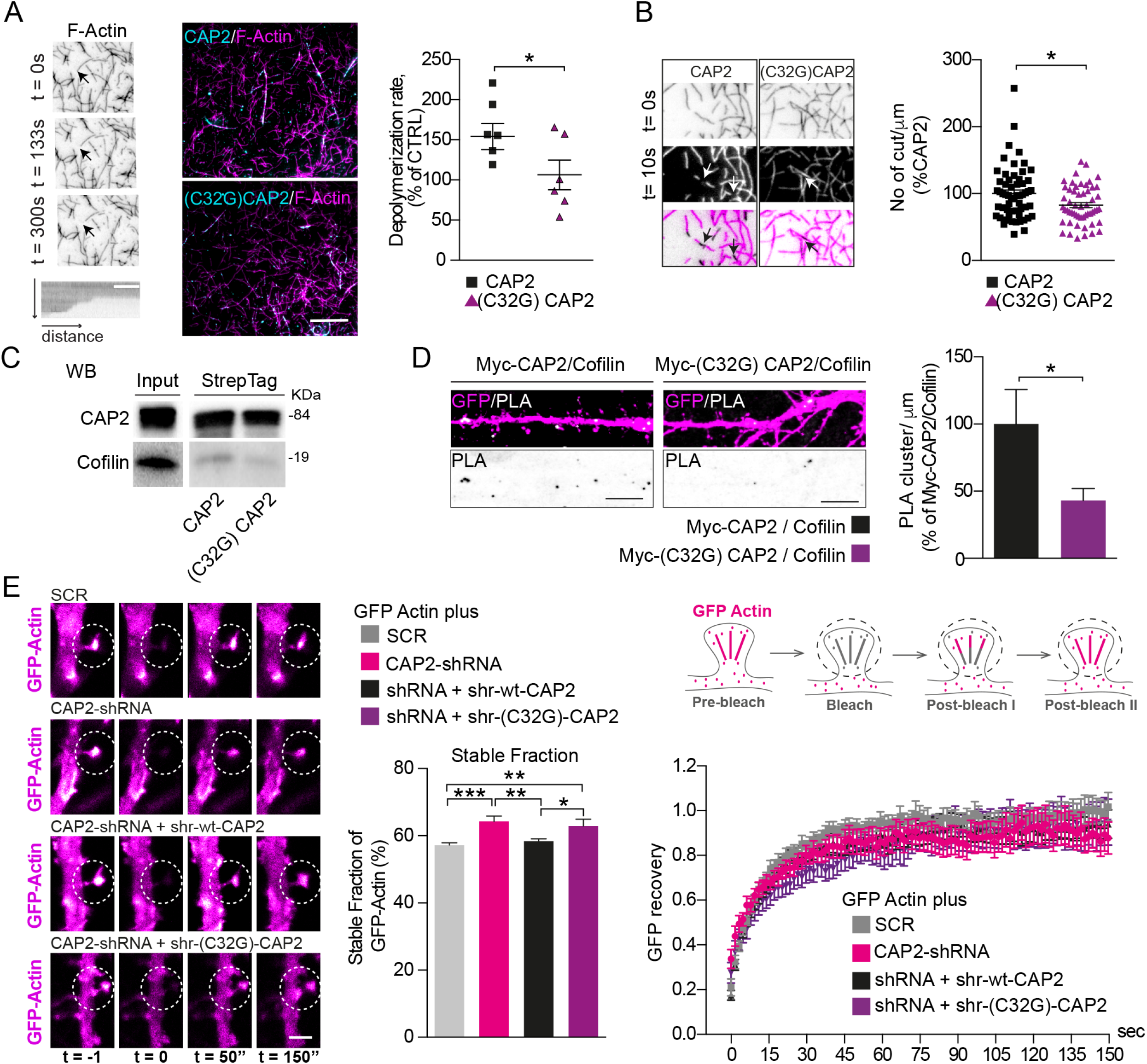
Cys^32^-dependent CAP2 dimerization is relevant for association with Cofilin and for controlling actin dynamics in spines. A Spontaneous *in vitro* depolymerization of F-actin-Alexa Fluor 568 visualized by TIRF microscopy. Left panels, actin filament undergoing depolymerization upon G-actin wash-out, arrowheads indicate the barbed ends of the filament. Right panels, overlay images show the actin filaments (*magenta*) and CAP2 and (C32G)CAP2 mutant (*cyan*). Scale bar = 5 µm. Kymographs analysis of F-actin depolymerization rate indicates that CAP2 increases the depolymerization rate of F-actin while the mutant (C32G)CAP2 abolished this effect (CAP2 vs (C32G)CAP2 \**P*=0.038, 2-tailed unpaired t-test, *n*=6 independent experiments). B Representative images of the analysis of the severing activity show an actin filament undergoing depolymerization in presence of either CAP2 or (C32G)CAP2 and arrowheads indicate the presence of cutting sites. The number of cuts/µm significantly decreases in presence of (C32G)CAP2 compared to CAP2 (CAP2 vs (C32G)CAP2, \**P*=0.01, 2-tailed unpaired t-test, *n*=58-54 filaments, each dot represents an individual filament) C WB of the Streptag CAP2 and (C32G)CAP2 fusion proteins shows that the mutation of C32 to G reduces CAP2 capability to bind Cofilin. D Images showing PLA signal between Cofilin and either Myc-CAP2 or Myc-(C32G)CAP2 (*white*) along MAP2 positive dendrites (*magenta*). Scale bar = 5 µm. Lower panels, inverted images of PLA signal (*black*). The mutation of Cys^32^ significantly reduces association with Cofilin (Myc-CAP2 vs Myc-(C32G)CAP2, \**P*=0.044, 2-tailed unpaired t-test, *n*=8-9 neurons from 2 independent experiments). E FRAP analysis of actin turnover in hippocampal neuron transfected with GFP actin and either CAP2-shRNA or CAP2-shRNA + shr-wt-CAP2 or CAP2-shRNA + shr-(C32G)-CAP2. Histogram represents the stable fraction of GFP-actin calculated for each individual time course of FRAP traces (SCR vs CAP2-shRNA \*\*\**P* = 0.0008; SCR vs CAP2-shRNA + shr-(C32G)-CAP2 \*\**P* = 0.007; CAP2-shRNA vs CAP2-shRNA + shr-wt-CAP2 \*\**P* = 0.004; CAP2-shRNA + shr-wt-CAP2 vs CAP2-shRNA + shr-(C32G)-CAP2 \**P* = 0.028, one-way ANOVA uncorrected Fisher’s LSD, *n*=13-15 neurons). Rightmost graph, FRAP curves (normalized to average pre-bleach fluorescence) were plotted from multiple single-spine ROIs for each condition, n = 13–15 neurons per condition, at least 3 spines for each neuron.

To confirm that the lack of Cys^32^ affects CAP2 association with Cofilin, PLA assays were performed in primary hippocampal neurons transfected with either Myc-CAP2 or Myc-(C32G)CAP2. The analysis of the PLA signals confirm a significant decrease in the association between Myc-(C32G)CAP2 and Cofilin when compared to the binding of Myc-CAP2 to Cofilin (Fig. 4D). For control experiments, when anti-Myc primary antibody or anti-cofilin antibody alone was used, no PLA signal was generated (EV4B). As control, we verified that the association of CAP2 with actin was not affected by the mutation of the Cys^32^ to Gly (EV4C). Therefore, the lack of the disulfide bond involving CAP2 Cys^32^ abolishes the CAP2 dimerization, reduces the Cofilin binding and, thereby, prevents the increase in actin depolymerization rate in TIRF experiments.

To directly address the functional consequences of the Cofilin-binding impairment of Myc-(C32G)CAP2 in the dendritic spines actin dynamics, we took advantage of fluorescence recovery after photobleaching (FRAP). The experiment was carried out in a single spine of hippocampal neuronal cells transfected with GFP-actin, that was partially incorporated into the actin filaments, and either CAP2-shRNA or its scrambled control sequence (Koskinen *et al*, 2012).

The recovery of GFP-actin fluorescence after spine photobleaching in CAP2 knockdown neurons was significantly attenuated (Fig. 4E) compared to control neurons. These data are consistent with a significant increase in the percentage of stable GFP-actin in spines lacking of CAP2 (Fig. 4E). To assess whether this effect could be specifically ascribed to CAP2 and to determine the role of the Cys^32^, we used shr-wt-CAP2 as a rescue and we generated and validated the mutant shr-wt-(C32G)CAP2 that was insensitive to the CAP2-shRNA (EV4D). As shown in Fig. 4E, the rise in the stable GFP-actin fraction caused by CAP2 knockdown was restored to control levels by the shr-wt-CAP2 but not by the mutant shr-(C32G)CAP2.

Taken together, these data suggest that the predominant effect of the loss of CAP2 on GFP-actin turnover is to reduce the pool of dynamic F-actin in spines, thus indicating that CAP2 is essential for actin turnover in spines. Most important, coexpression of (C32G)CAP2 mutant could not rescue the FRAP defects, indicating that dimer formation and, thereby, Cofilin association are critical for the regulation of actin dynamics in spines.

### Long-term potentiation triggers CAP2 dimer formation

Given the key role of Cofilin-mediated actin dynamics in the structural remodeling of spines during activity-dependent synaptic plasticity phenomena (Bosch *et al*, 2014), we tested whether LTP and LTD modulate CAP2 dimerization, and consequently its association with Cofilin.

First, we investigated CAP2 homodimer localization in the postsynaptic compartment. Western Blot analyses showed that CAP2 dimer levels are higher in the postsynaptic TIF compared to the total homogenate (EV5A) in both rat and human hippocampus, thus confirming the existence of CAP2 dimer in human and revealing the enrichment of the CAP2 dimer in this subcellular compartment.

Second, we verified whether LTP regulates CAP2 synaptic availability in the dendritic spines. To this, we induced cLTP in hippocampal cultures that results in prolonged NMDA receptor-dependent LTP (Marcello *et al*, 2013). 30 min after cLTP induction, we purified TIF from control and cLTP-treated hippocampal neurons. Western blot analysis revealed that cLTP elicited an increase in the CAP2 levels in the TIF fraction compared to control neurons, without affecting the total levels of the protein (Fig. 5A). Notably, cLTP stimulation caused also a significant increase in CAP2 dimer levels in TIF (Fig. 5B). Secondly, to evaluate whether these effects could be ascribed to the CAP2 Cys^32^-dependent dimerization, we analyzed the PLA signal upon cLTP induction in hippocampal neurons transfected with EGFP-CAP2 and either Myc-CAP2 or Myc-(C32G)CAP2. The cLTP induction significantly increased the PLA signal in neurons transfected with EGFP-CAP2 and Myc-CAP2, confirming the LTP-triggered augment in CAP2 self-association. On the other hand, no modifications in PLA signals upon LTP induction were detected when EGFP-CAP2 and Myc-(C32G)CAP2 were expressed (Fig. 5C). This effect is specifically related to LTP, since both PLA and Western Blot analysis of the synaptic fraction revealed no variations in CAP2 synaptic levels and in dimer formation after cLTD induction (EV5 B, C, D).

**Figure 5.**
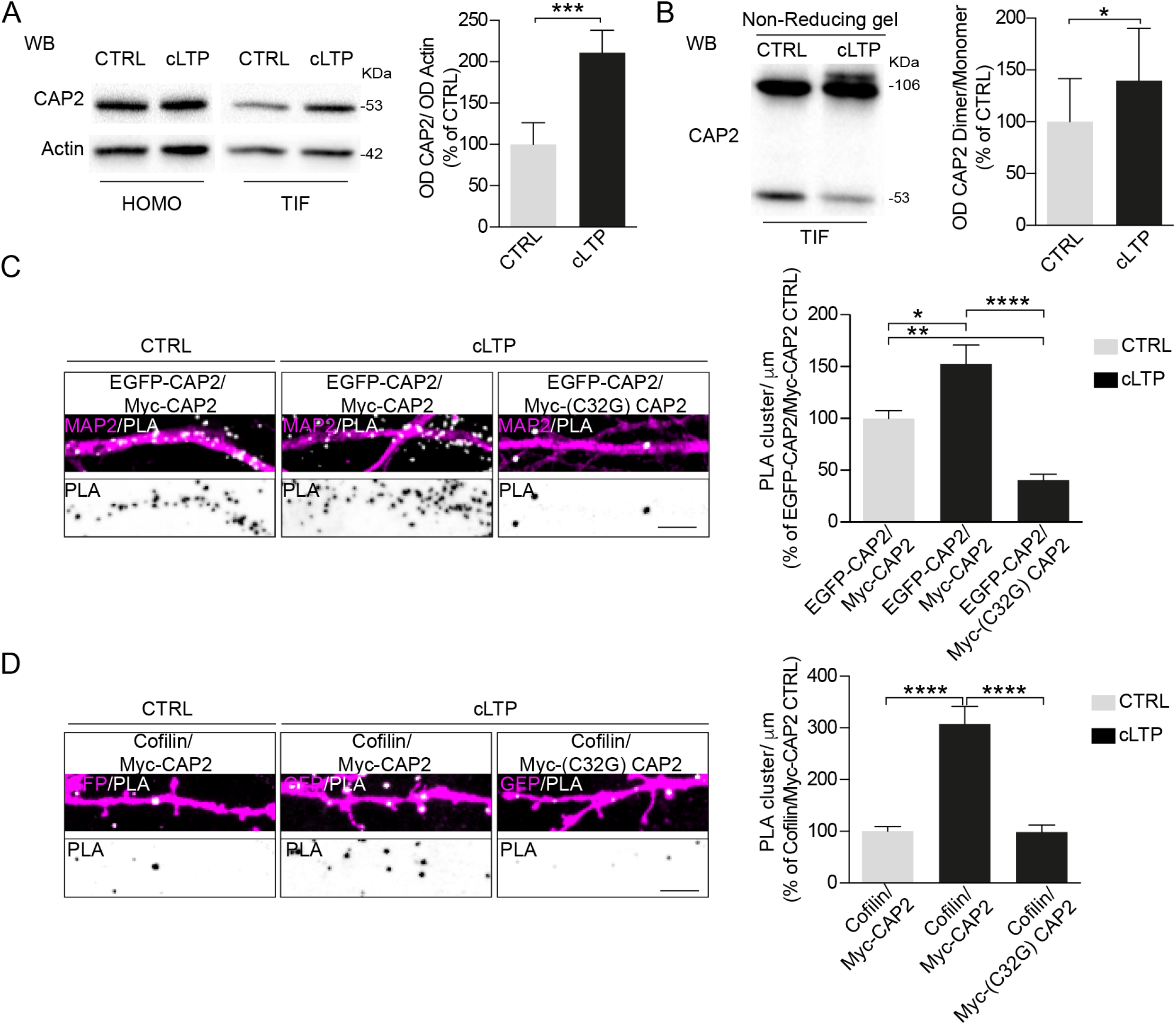
LTP promotes CAP2 synaptic localization, dimer formation and association with Cofilin. A Representative WB of Homo and TIF obtained from control and cLTP-treated neurons, 30 min after the cLTP induction. cLTP promotes CAP2 enrichment in the TIF (CTRL vs cLTP \*\*\**P*=0.0008, 2-tailed paired t-test*, n*=5 independent experiments). B cLTP increases CAP2 dimer/monomer ratio in the TIF fraction of cLTP-treated neurons (CTRL vs cLTP, \**P*=0.042, 2-tailed paired t-test, *n*=5 independent experiments). C PLA representative images showing the proximity between EGFP-CAP2 and either Myc-CAP2 or Myc-(C32G)CAP2 (*white*) along MAP2 positive dendrites (*magenta*) 30 min after cLTP induction. Lower panels, inverted images of PLA signal (*black*), scale bar = 5 µm (EGFP-CAP2/Myc-CAP2 CTRL vs EGFP-CAP2/Myc-CAP2 cLTP \**P*=0.01, EGFP-CAP2/Myc-CAP2 CTRL vs EGFP-CAP2/Myc-(C32G)CAP2 cLTP \*\**P*=0.008; EGFP-CAP2/Myc-CAP2 cLTP vs EGFP-CAP2/Myc-(C32G)CAP2 cLTP **** *P* < 0.0001, one-way Anova, Bonferroni’s multiple comparisons test, *n*=6-8 neurons per condition from at least 2 independent experiments). D PLA images revealing the closeness between Cofilin and either Myc-CAP2 or Myc-(C32G)CAP2 (*white*) along dendrites of GFP transfected hippocampal neurons (*magenta*) after cLTP induction. Lower panels, inverted images of PLA signal (*black*), scale bar = 5 µm, MycCAP2/Cofilin CTRL vs MycCAP2/Cofilin cLTP \*\*\*\**P* < 0.0001; MycCAP2/Cofilin cLTP vs Myc-(C32G)-CAP2/Cofilin cLTP \*\*\*\**P* < 0.0001; one-way Anova, Bonferroni’s multiple comparisons test, *n*= 5-6 neurons per condition from at least 2 independent experiments).

To determine if the increase in LTP-driven CAP2 dimer formation affects the association with Cofilin, we performed PLA assays after cLTP induction in neurons transfected with either Myc-CAP2 or Myc-(C32G)CAP2. As shown in Fig. 5D, LTP fosters the association of Cofilin with CAP2. Likewise, we detected a significant increase in the association between endogenous CAP2 and Cofilin upon LTP induction, indicating that this effect was not related to the overexpression of Myc-CAP2 construct (EV5E). The Cys^32^-dependent self-association of CAP2 is required because the Myc-(C32G)CAP2 mutant prevents the LTP-induced increase in CAP2/Cofilin interaction (Fig. 5D).

Overall, these findings suggest that LTP triggers CAP2 translocation to the spine and the formation of Cys^32^-dependent dimers, thus promoting the association of CAP2 with Cofilin.

### CAP2 dimer formation is critical for LTP

In the initial phase of LTP the actin cytoskeleton is rapidly remodeled while active Cofilin is massively transported to the spine (Bosch *et al*, 2014). In light of this observation and considering that LTP elicits CAP2 dimer formation in the postsynaptic compartment, thus promoting CAP2 association with Cofilin, we attempted to determine whether the hampering CAP2 dimerization impairs LTP-driven enrichment of Cofilin in the spine. To this, we induced cLTP in primary hippocampal neurons transfected with either CAP2-shRNA or the corresponding control scramble sequence. These constructs express the fluorescent protein mCherry under the control of the neuron-specific CaMKII promoter. To assess the magnitude of accumulation, we took advantage of structured illumination microscopy (SIM) to calculate the relative concentration of Cofilin in the spine by dividing Cofilin intensity by mCherry intensity (Bosch *et al*, 2014). As previously demonstrated (Bosch *et al*, 2014), after 30 min from cLTP induction, Cofilin is highly enriched in the spines of neurons transfected with the scramble sequence, while the down-regulation of CAP2 prevents the LTP-triggered translocation of Cofilin in the spine. Importantly, the transfection of the shRNA-resistant CAP2 construct (shr-wt-CAP2), but not the expression of the shRNA-resistant (C32G)CAP2 mutant [shr-(C32G)-CAP2], restores the LTP-driven translocation of Cofilin to the spines (Fig. 6A). These data demonstrate that CAP2 and the Cys^32^-dependent dimerization of the protein are essential for the LTP-triggered Cofilin trafficking to spines.

**Fig. 6.**
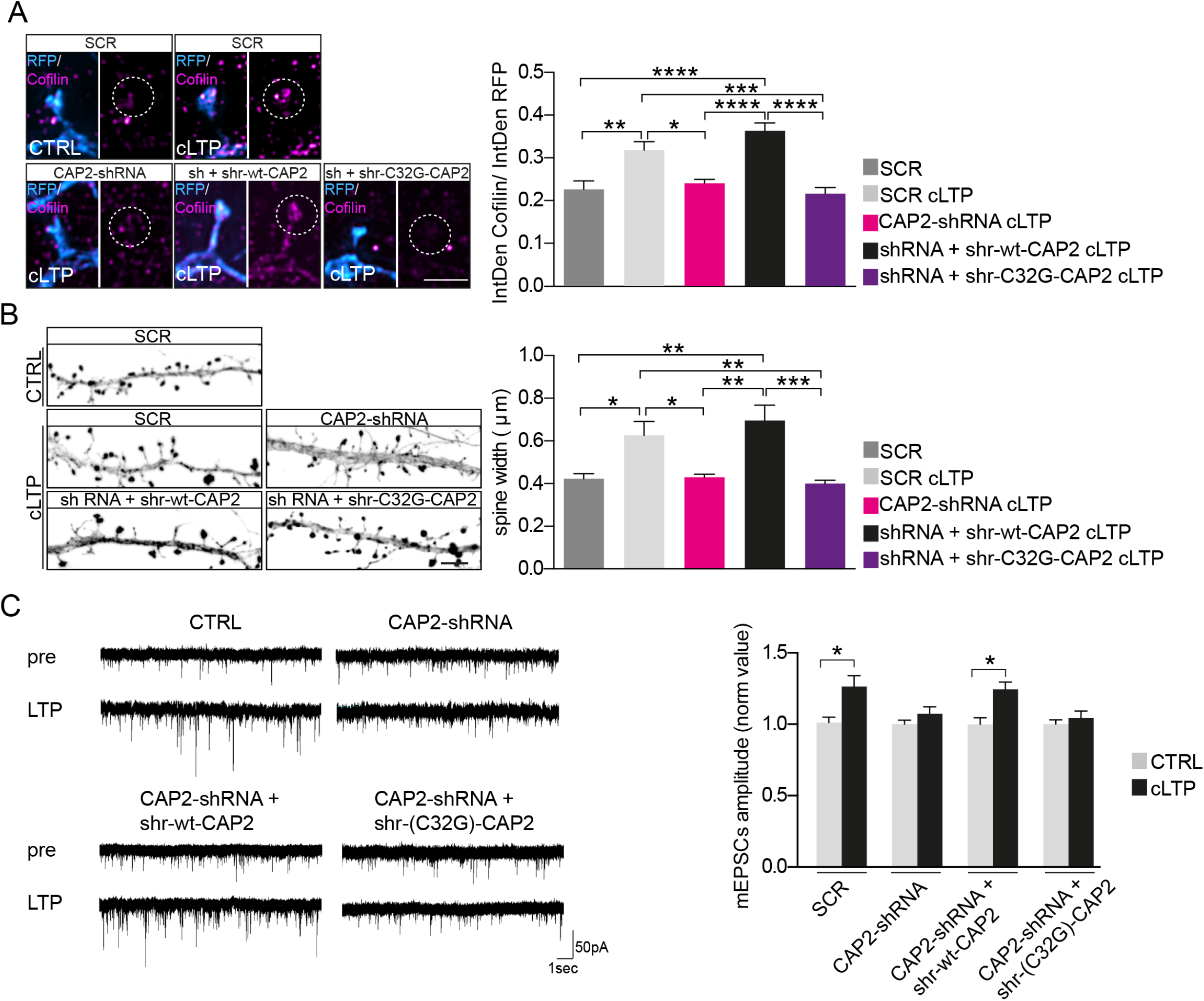
Cys^32^-dependent CAP2 dimer is required for Cofilin translocation upon LTP and for structural plasticity changes and for synaptic transmission potentiation. A SIM images of Cofilin and RFP staining of primary hippocampal neurons transfected with vectors expressing RFP and either scramble sequence (SCR) or CAP2-shRNA or CAP2-shRNA + shr-wt-CAP2 or CAP2-shRNA + shr-(C32G)-CAP2, 30 min after cLTP induction. Scale bar = 2 µm. Histograms show the quantification of the ratio between the integrated density (IntDen) of Cofilin and RFP staining in dendritic spines (SCR vs SCR cLTP ** *P* = 0.0012; SCR cLTP vs CAP2-shRNA cLTP * *P* = 0.002; SCR vs CAP2-shRNA + shr-wt-CAP2 cLTP **** *P* < 0.0001; SCR cLTP vs CAP2-shRNA + shr-(C32G)-CAP2 cLTP *** *P*= 0.0004; CAP2-shRNA cLTP vs CAP2-shRNA + shr-wt-CAP2 cLTP **** *P* < 0.0001; CAP2-shRNA + shr-wt-CAP2 cLTP vs CAP2-shRNA + shr-(C32G)-CAP2 cLTP **** *P* < 0.0001,, one-way Anova, Bonferroni’s multiple comparisons test, *n*=7-8 neurons per condition from at least 2 independent experiments). B Confocal images of RFP staining of primary hippocampal neurons transfected with vectors expressing RFP and either SCR or CAP2-shRNA or CAP2-shRNA + shr-wt-CAP2 or CAP2-shRNA + shr-(C32G)-CAP2, 30 min after cLTP induction; scale bar = 5 µm. Histograms show the quantification of dendritic spine head width (SCR vs SCR cLTP \**P* = 0.033; SCR cLTP vs CAP2-shRNA cLTP \**P* = 0.037; SCR vs CAP2-shRNA + shr-wt-CAP2 cLTP ** *P* = 0.001; SCR cLTP vs CAP2-shRNA + shr-(C32G)-CAP2 cLTP \*\**P* = 0.009; CAP2-shRNA cLTP vs CAP2-shRNA + shr-wt-CAP2 cLTP \*\**P* = 0.0014; CAP2-shRNA + shr-wt-CAP2 cLTP vs CAP2-shRNA + shr-(C32G)-CAP2 cLTP *** *P* =0.0003, one-way Anova, Bonferroni’s multiple comparisons test, *n*=10-11 neurons per condition from at least 2 independent experiments). C Electrophysiological recordings of mEPSCs before and after chemical LTP (cLTP) induction in 15 DIV hippocampal neurons. (Left) Representative mEPSC traces. Differently from CTRL and shr-wt-CAP2 cells, CAP2-shRNA and CAP2-shRNA+shr-(C32G)-CAP2 neurons are unable to undergo cLTP. mEPSCs amplitude (normalized values): CTRL= 1±0.03 (n=18) vs cLTP: 1.26±0.07 (n=22); CAP2-shRNA: 1±0.02 (n=15) vs cLTP: 1± 0.02 (n=20); shr-wt-CAP2: 1±0.04 (n=19) vs cLTP:1.25±0.05 (n=23); shRNA+shr-(C32G)-CAP2: 1±0.031 (n=18) vs cLTP: 1± 0.04 (n=26). One Way Anova, Holm-Sidak’s multiple comparisons test, *P*<0.0001.

Since Cofilin translocation to spines during LTP is required for actin cytoskeleton dynamics and spine enlargement, we verified whether the loss of Cys^32^ and, thereby, of CAP2 dimerization impairs LTP-induced spine remodeling. An accurate analysis of dendritic spine morphology demonstrated that cLTP induces the expected spine head enlargement in control conditions, but not in neurons transfected with CAP2-shRNA (Fig. 6B). Interestingly, LTP-dependent spine enlargement could be rescued by shr-wt-CAP2, but not by the mutant shr-(C32G)-CAP2 (Fig. 6B).

Furthermore, taking advantage of electrophysiological recordings of mEPSCs, we addressed the functional impact of CAP2 down-regulation on hippocampal plasticity by induction of cLTP. As indicated in Fig. 6C, cLTP mediates a significant increment in mEPSCs amplitude of excitatory synaptic events only in control cultures whereas no potentiation occurs in CAP2-shRNA neurons. Similarly, cultures transfected with the shr-(C32G)-CAP2 mutants do not display any potentiation that is rescued only in the shRNA+shr-wt-CAP2 control group. Taken together, these results indicate that synaptic plasticity is affected by CAP2 and that mutation in the residue Cys^32^ is sufficient for defective CAP2 activity and LTP induction.

### CAP2 dimer association with Cofilin is altered in AD patients

Since Cys^32^-dependent CAP2 homodimerization is crucial for the regulation of dendritic spine morphology and synaptic plasticity via Cofilin binding, we wondered whether alterations of this mechanism were detectable in AD synapses, where we found an aberrant increased level of Cofilin along with a significant reduction in CAP2 (Fig.1).

The analysis of the CAP2 dimer/monomer ratio in APP/PS1 and wild-type mice hippocampi revealed a significant decrease of dimer levels in the postsynaptic fraction, but not in the total homogenate, of APP/PS1 mice when compared to wild-type mice (Fig. 7A). We examined the CAP2 homodimer formation in AD patients’ hippocampi and the quantitative results showed a significant reduction in CAP2 dimer/monomer ratio specifically in the synapse of AD patients compared to HC, while no changes were detected in the total homogenate (Fig. 7B)

**Fig. 7.**
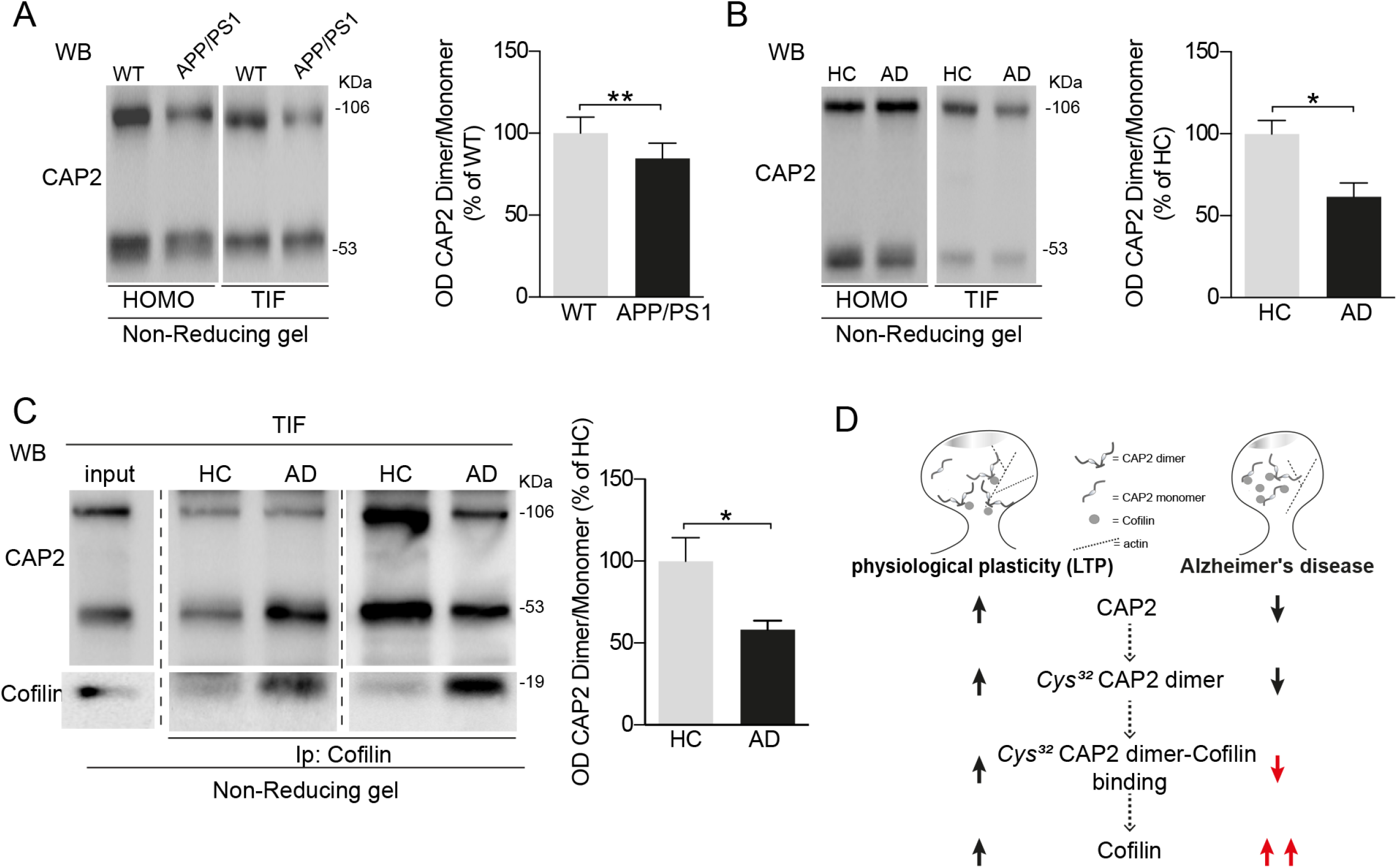
Cys^32^-dependent CAP2 dimer levels and CAP2 association to Cofilin are affected in AD synapses. A. Representative WB of CAP2 dimer level in the HOMO and TIF hippocampal fraction of APP/PS1 vs WT animals. Quantitative analysis shows the decreased ratio of OD. CAP2 dimer on the OD CAP2 monomer of APP/PS1 compeared to controls (TIF, WT vs APP/PS1 \*\**P*= 0.003, 2-tailed paired t-test, *n*=6). B Representative WB showing the levels of CAP2 monomer and dimer in the HOMO and TIF samples of AD and HC hippocampal samples. Quantitative analysis shows that CAP2 dimer/monomer ratio is reduced in AD patients TIF but not in HOMO (TIF, HC vs AD \**P*= 0.014, 2-tailed paired t-test, *n*=8). C TIF from hippocampi of HC and AD patients were Ip with anti-Cofilin antibody and CAP2 coprecipitation evaluated in non-denaturating condition. Samples derive from the same experiment and gels/blots were processed in parallel. Cofilin binds preferentially the CAP2 monomer in TIF fraction of AD compared with HC (CAP2 dimer/monomer, HC vs AD \**P* = 0.023, 2-tailed unpaired t-test, *n*=8). D A working model for the involvement of CAP2 in the pathogenesis of AD. LTP triggers CAP2 translocation to spines, the formation of Cys^32^-dependent CAP2 dimers and binding to Cofilin, thus leading to an increase in Cofilin synaptic localization. In AD patients we observed a reduction in CAP2 expression and synaptic localization leading to a decrease formation of Cys^32^-dependent dimers, an aberrant association of Cofilin to CAP2 monomers and a pathological increased localization of Cofilin into spines.

Since CAP2 dimer formation is relevant for CAP2 association with Cofilin, we verified whether Cofilin binding to CAP2 monomer and dimer is affected in AD. Given that Cofilin is aberrantly increased in the AD hippocampi postsynapse, we focused on TIF samples obtained from HC and AD patients. TIF samples were immunoprecipitated with Cofilin antibody and loaded onto a non-reducing gel. Western Blot analysis carried out with a CAP2 antibody revealed that Cofilin specifically precipitates both CAP2 monomer and dimer in human postsynaptic compartment (EV6), confirming CAP2 dimer and monomer as binding partners of Cofilin in human. The quantitative analysis of coimmunoprecipitation experiments revealed that the ratio of CAP2 dimer/monomer bound to Cofilin is significantly reduced in AD patients when compared to HC (Fig. 7C). These results indicate that Cofilin preferentially associates with CAP2 monomer than with the CAP2 dimer in AD postsynaptic compartment.

In light of the above, we tested whether the association between single nucleotide polymorphisms of CAP2 Cys^32^ (rs750613178 variant) and AD might occur. To this aim, a preliminary screening experiment of genetic polymorphism has been performed on 70 AD patients (age=68.2±16.3; female%= 76.8). No genetic variations of CAP2 Cys^32^ were detected in AD patients, suggesting that other cellular mechanisms might be responsible for the decreased synaptic dimer levels in AD.

## Discussion

A major question in Alzheimer disease pathogenesis is to define the mechanism guiding alteration of synaptic plasticity and spine dysmorphogenesis. Here, we show that CAP2 is able to control Cofilin synaptic availability during activity-dependent plasticity phenomena (Fig. 7D) and it is essential for actin turnover. Remarkably, this mechanism is specifically altered in the hippocampus of AD patients and of an AD mouse model (Fig. 7D).

CAP2 has been reported as a multi-domain actin binding protein, expressed in specific tissues, including the brain (Peche *et al*, 2007; Kumar *et al*, 2016). Our study provides a detailed characterization of CAP2 in neurons bringing into focus conclusive evidences for both CAP2 localization in the postsynaptic compartment and its pivotal role in synaptic plasticity phenomena. This has been demonstrated with different approaches. First, STED nanoscopy and biochemical fractionation approaches showed that CAP2 is localized in the postsynaptic compartment, in the region just underneath the PSD where F-actin is enriched (Korobova & Svitkina, 2010). Secondly, we observed that the down-regulation of CAP2 impairs neuronal architecture and spine shape, together with a decrease in synaptic excitatory transmission. Such alterations are translated into an impairment of plasticity phenomena.

Our results point to a defect in actin dynamics in spines, shown by FRAP experiments, as the potential mechanism underlying such damaged neuronal phenotype. In fact, the analysis of FRAP experiments revealed that the down-regulation of CAP2 results in reduced actin turnover and, thereby, in a slight but significant increase in the actin stable pool in spines. Along these lines, CAP2 can be defined as an actin cytoskeleton modifier, essential for normal actin turnover in spines. In accordance with this hypothesis and considering that actin polymerization in spines is required for functional LTP (Okamoto *et al*, 2004), we observed that the down-regulation of CAP2 leads to the lack of both potentiation and spine head enlargement expected upon cLTP induction.

Why is CAP2 critical for plasticity phenomena? CAP2 has been reported as a binding partner of Cofilin (Kumar *et al*, 2016), a master regulator of postsynaptic actin dynamics and structural plasticity (Rust, 2015a). Upon LTP induction, Cofilin activity is first increased and subsequently decreased, allowing first for actin remodeling and then for an increase in F-actin and associated spine enlargement (Fukazawa *et al*, 2003; Chen *et al*, 2007; Gu *et al*, 2010). Even though Cofilin phosphorylation is considered one of the major mechanisms controlling its activity (Van Troys *et al*, 2008; Bernstein & Bamburg, 2010; Rust, 2015a), the regulation of its availability and retention in spines could also contribute to Cofilin-dependent sculpturing of the cytoskeleton. Cofilin indeed accumulates in regions of the spine that contain a dynamic F-actin network responsible for spine morphological changes during synaptic plasticity (Racz & Weinberg, 2006). Importantly, Cofilin is massively translocated to the spine upon LTP induction (Bosch *et al*, 2014) and our findings show that CAP2 down-regulation blocks the LTP-triggered enrichment of Cofilin in spines. The impaired Cofilin localization in spines and thereby structural and functional synaptic plasticity could be rescued by cotransfection of the cells with an shRNA-resistant CAP2 construct, confirming that the described CAP2-knockdown phenotypes indeed resulted from diminished CAP2 expression.

Furthermore, we also addressed the key question regarding the molecular determinants relevant for CAP2/Cofilin association. A careful mapping of CAP2 structure was carried out and, by using different approaches, we showed that the N-terminal domain of CAP2 is responsible for its self-association. In particular, in this region we identified the Cys^32^ as the amino acid involved in the formation of disulfide cross-linked CAP2 dimers. The mutation of Cys^32^ to Gly dramatically reduces, but not completely abolishes, CAP2 self-association and the binding to Cofilin, suggesting that CAP2 can still form aggregates capable of interacting with Cofilin. However, the lack of Cys^32^-dependent CAP2 homodimer results in a loss of function of CAP2/Cofilin complex on actin turnover, highlighting the importance of such disulfide bond for the operational role of CAP2 in the synapse. In fact, the CAP2 construct carrying the mutation Cys^32^ to Gly significantly attenuates the capability of CAP2 of promoting actin filaments depolymerization in an *in vitro* assay. In addition, in a cellular context, FRAP experiments demonstrated that the CAP2 mutant lacking Cys^32^ fails to rescue the abnormal treadmilling of actin in the spines of CAP2-knockdown neurons. A similar molecular pathway has been shown for CAP1, since Cys^29^-dependent homodimerization of CAP1 affects binding to Cofilin and F-actin stability (Liu *et al*, 2018). Notably, the Cys^29^ residue of CAP1 is not conserved in human, thus limiting the relevance of CAP1-dependent mechanism and highlighting the role of CAP2 as main regulator of Cofilin in human neuronal cells.

To strengthen the critical role of CAP2 homodimer in the spine, we show that it is localized in the postsynaptic compartment and it undergoes a dynamic regulation of its synaptic levels by activity-dependent synaptic plasticity. The LTP-induced increase in CAP2 localization in spines promotes the formation of CAP2 dimers through Cys^32^, and thereby fosters CAP2 association with Cofilin. This event is required for LTP-induced translocation of Cofilin to the spine and for structural changes and the potentiation of synaptic transmission underlying LTP expression. Indeed, the CAP2 construct carrying the mutation Cys^32^ to Gly, incapable of dimerization, is not able to rescue the LTP and the LTP-triggered enlargement of spine head in CAP2-knockdown neurons, as the wild-type CAP2 construct. In addition, the SIM analysis of Cofilin localization in spines revealed that the CAP2 mutant failed to restore the expected enrichment of Cofilin in spines upon LTP. These data suggest that the increased synaptic availability of CAP2 monomers in the synapse triggers the dimer formation and the downstream events, even though we can’t exclude that post-translational modifications or interacting proteins could affect this pathway. Of note, the induction of LTD doesn’t affect CAP2 localization in the spine and dimer formation, highlighting the specificity of this mechanism for LTP.

Cofilin has been implicated in the pathophysiology of AD, since Cofilin accumulates in senile plaques in AD tissue and AD mouse models (Bamburg & Bernstein, 2016). However, the mechanisms by which Cofilin contributes to AD pathogenesis remain quite controversial, as either an excessive activation (Kim *et al*, 2013) or inactivation (Barone *et al*, 2014; Rush *et al*, 2018) have been reported in AD patients. Here we changed perspective, and bring into focus Cofilin localization in spines in AD, because it is directly regulated by LTP, as well as its association with CAP2 (Bosch *et al*, 2014). The CAP2 dimer-dependent mechanism that regulates Cofilin availability at the synapse is specifically affected in the hippocampus, but not in the SFG, of both AD patients and APP/PS1 mice. In the hippocampal synapses of AD patients and AD mouse model, we measured a dramatic increase in Cofilin levels, along with a reduction in CAP2 synaptic availability, and accordingly a decrease in CAP2 dimer formation at the synapse. Furthermore, we noticed that in AD patients’ hippocampi Cofilin preferentially associates with the CAP2 monomer than to the CAP2 dimer, suggesting the presence of an ineffective CAP2/Cofilin complex in AD hippocampal synapses that could contribute to the loss of structural plasticity of spines in AD.

Together, our findings describe a novel CAP2-dependent mechanism controlling Cofilin synaptic availability and actin turnover in spines, which is closely interdependent to LTP. Furthermore, we show that the main players of this pathway are altered in AD, adding new pieces to the puzzle in the understanding of the complex and coordinated events leading to early synaptic dysfunction and plasticity alterations in AD pathogenesis.

## Materials and methods

### Human brain tissue

The hippocampus and SFG samples of AD patients and of age-matched healthy controls (HC) were obtained from the Netherlands Brain Bank (NBB). Established Braak and Braak criteria were used to categorize AD tissues (Braak & Braak, 1991). AD patients fulfilled Braak 4 and 5 stages. Accordingly, in AD cases, there were tangles and neuritic plaques in hippocampus. HC had no history of psychiatric or neurological disease and no evidence of age-related neurodegeneration. Detailed information is reported in table 1 and 2.

### Triton-Insoluble synaptic membrane preparation

Triton insoluble fraction (TIF), a fraction highly enriched in all categories of postsynaptic density proteins (i.e., receptor, signaling, scaffolding, and cytoskeletal elements) absent of presynaptic markers, was obtained from human hippocampus and SFG specimens and APP/PS1 and wild-type hippocampal samples as previously described (Marcello *et al*, 2012b; Epis *et al*, 2010). In order to avoid protein degradation, AD samples were paired to HC samples and processed at the same time. The procedure was performed at least twice to have two experimental replicates. The TIF was purified from primary hippocampal cultures as previously described (Marcello *et al*, 2007).

### Treatments of neuronal cultures

To induce chemical LTP (cLTP), hippocampal neuronal cultures were first incubated in artificial cerebrospinal fluid (ACSF) for 30 min: 125 mM NaCl, 2.5 mM KCl, 1 mM MgCl_2_, 2 mM CaCl_2_, 33 mM D-glucose, and 25 mM HEPES (pH 7.3; 320 mosM final), followed by stimulation with 50 µM forskolin, 0.1 µM rolipram, and 100 µM picrotoxin (Tocris) in ACSF without MgCl_2_. After 16 min of stimulation, neurons were replaced in regular ACSF for 15 min and, after treatment, samples were processed (Marcello *et al*, 2013). To induce chemical LTD (cLTD), neuronal cultures were incubated in ACSF for 30 min, followed by stimulation with 50 µM NMDA (Sigma-Aldrich) in ACSF. After 10 min of stimulation, neurons were replaced in regular ACSF for 20 min and then subjected to the biochemical and imaging studies (Marcello *et al*, 2013). Stimulation of synaptic NMDA receptors (SS) was obtained by treating hippocampal neurons with 50 mM Bicuculline (Tocris), 2.5 mM 4-AP and 5 mM ifenprodil in Neurobasal medium supplemented with B27 (Dinamarca *et al*, 2016).

### Co-immunoprecipitation assays

Aliquots of 20 µg of TIF obtained from human hippocampus were incubated with an antibody (Ab) against Cofilin overnight at 4°C in a final volume of 150 µL of RIA buffer [200 mM NaCl, 10 mM ethylene-diaminetetra-acetic acid (EDTA), 10 mM Na_2_HPO_4_, 0.5% NP-40, 0.1% sodium dodecyl sulfate (SDS)]. SureBeads Protein A/G Magnetic Beads (Bio-Rad) were used to precipitate the immunocomplex, the beads were resuspended in sample buffer without β-mercaptoethanol and heated for 3 min before the loading onto SDS-PAGE. Beads were collected by centrifugation and applied onto SDS-PAGE; the precipitated immunocomplex was revealed by anti-Cofilin and anti-CAP2 antibody. Heterologous co-immunoprecipitation experiments were carried out from lysates of either COS-7 or HEK293 cells transfected with different combinations of Myc-CAP2, EGFP-CAP2 or EGFP tagged truncated constructs of CAP2. Cells were harvested and proteins extracted as previously described (Marcello *et al*, 2010). The same immunoprecipitation protocol was used, incubating aliquots of 100 µg of HEK293/COS-7 lysates with an anti-Myc antibody. Protein A/G Agarose beads (Pierce) were added, and incubation was continued for 2 h at room temperature (RT) on the rotator mixer. Beads were collected and washed with RIA buffer for 3 times. Sample buffer for SDS-PAGE was added and the mixture was heated for 10 min. The precipitated immunocomplex was revealed with either anti-GFP or anti-Myc or anti-actin antibodies.

### In situ proximity ligation assay (PLA)

PLA was performed in primary neuronal cultures as previously described (Dinamarca *et al*, 2016). Primary hippocampal neurons were fixed with 4% Paraformaldehyde (PFA) - 4% sucrose for 5 min at 4°C and washed several times with PBS. Neurons were permeabilized with 0.1% Triton X-100 in PBS for 15 min at RT. After incubation with the blocking solution of the PLA kit (Duolink® PLA Technology), cells were incubated overnight with the primary antibodies at 4°C. According to the manufacturer’s instructions, after several washing with solution A, secondary probes attached to oligonucleotides were added and the oligonucleotides of the bound probes where ligated and amplified by a fluorescent polymerase that visualizes the PLA signal. To stain MAP2 or the transfected GFP, cells were washed with PBS for 30 min and then the immunocytochemistry protocol was used. Cells were mounted on slides in Fluoromount™ Aqueous Mounting Medium (Sigma-Aldrich).

### Cell culture electrophysiology

Whole cell voltage-clamp recordings were performed on rat hippocampal neurons transfected at day *in vitro* (DIV) 10 and maintained in culture for 15-16 DIV. Recording pipettes were fabricated from borosilicate glass capillary with a tip resistance of 3–5 MΩ and filled with an intracellular solution of the following composition (in mM): 130 potassium gluconate, 10 KCl, 1 EGTA, 10 Hepes, 2 MgCl2, 4 MgATP, and 0.3 Tris-GTP. During recordings of miniature excitatory postsynaptic currents (mEPSCs) cells were bathed in a standard external solution containing (in mM): 125 NaCl, 5 KCl, 1.2 MgSO4, 1.2 KH2PO4, 2 CaCl2, 6 glucose and 25 HEPES-NaOH, pH 7.4, with also tetrodotoxin (TTX-1µM), bicuculline (20µM) and strychnine (1µM). For chemical LTP experiments, recordings of mEPSCs were performed applying glycine (100µM) for 3min at room temperature in Mg^2+^-free KRH, also containing TTX, bicuculline and strychnine. Recordings were performed at room temperature in voltage clamp mode using a Multiclamp 700B amplifier (Molecular Devices) and pClamp-10 software (Axon Instruments). Series resistance ranged from 10 MΩ to 20 MΩ and was monitored for consistency during recordings. Cells in culture with leak currents >200pA were excluded from the analysis. Signals were amplified, sampled at 10kHz, filtered to 2 or 3KHz and analyzed using pClamp 10 data acquisition and analysis program.

### Ethical approvals

All procedures performed in studies involving human participants were in accordance with the ethical standards of the institutional research committee (Brescia Ethics Committee, protocol #NP2806) and with the 1964 Helsinki declaration and its later amendments or comparable ethical standards.

All procedures performed in studies involving animals were in accordance with the ethical standards of the Institutional Animal Care and Use Committee of University of Milan (Italian Ministry of Health permit #326/2015, #5247B.N.YCK/2018, #295/2012-A, #497/2015-PR) and of the internal animal welfare authorities of the University of Marburg (references: AK-6-2014-Rust), at which the studies were conducted.

## Acknowledgements

We thank A. Longhi and E. Zianni for technical assistance and Alessia Mariani, Filippo La Greca, Ramona Stringhi and Michael Wolf for excellent practical work. This project has received funding from the European Union’s Horizon 2020 research and innovation program under the Marie Skłodowska-Curie grant agreement No 676144 (Synaptic Dysfunction in Alzheimer Disease, SyDAD) to MDL, from the Italian Ministry of University and Research (PRIN 2015N4FKJ4 to MDL, Fondo per il Finanziamento delle Attività Base di Ricerca FFABR18_10 to EM, MIUR Progetto Eccellenza), from Fondazione Cariplo to EM (Grant n. 2018 - 0511), from AIRAlzh Onlus-COOP Italia (fellowship to SP), from an intramural grant of University of Milan to EM (Fondo di sviluppo unimi-linea2 - PSR2015-1716GRACA02_05_M and PSR2017_DIP_022_03), from the Deutsche Forschungsgemeinschaft (DFG) [Emmy-Noether Programm (MI 1923/1-1) and FOR2419 (MI 1923/2-1 and MI 1923/2-2)] to MM and from DFG (SFB1158, project A08) to DM. This work was supported by MIUR - PON “Ricerca e Innovazione” PerMedNet project (ARS01_01226). We thank UMIF (UKE, Hamburg) for access to their STED imaging system. The brain tissues were obtained from the Netherlands Brain Bank, Netherlands Institute for Neuroscience, Amsterdam.

## Authors contributions

SP and LV carried out and analyzed the biochemical and imaging experiments. LP and FA performed electrophysiology experiments and their analysis. BA and DM designed and carried out quantitative real-time PCR, JY produced the recombinant adeno-associated viruses, AK and MM contributed to TIRF analysis and STED nanoscopy, EB and BB analyzed SNPs in AD patients, DDM performed the multiple-sequence alignment, MR contributed to the analysis of CAP2 during development. EM, SP, LV and MDL conceived the study and wrote the manuscript. FG, DM, FA, DDM, MR, MM contributed to the writing. All authors read and approved the final manuscript.

## The paper explained

### Problem

Alterations of synaptic plasticity and spines remodeling is a key aspect in Alzheimer disease pathogenesis. Cofilin is one of the major regulators of actin dynamics in spines where it is required for structural synaptic plasticity, and its alterations have been reported in Alzheimer disease. However, a clear picture of underlying mechanisms is still poorly understood making it difficult to clearly identify possible targets for future therapeutic development.

### Results

We identify a new regulator of cofilin - CAP2 - which governs the levels of cofilin in spines. By dimer formation, CAP2 binds to cofilin and controls normal actin turnover in spines. We show that a Cys in position 32 within CAP2 sequence is essential for dimerization.

We demonstrated that long-term potentiation (but not long-term depression) is promoting dimer formation and binding to cofilin, promoting cofilin translocation into spines.

Importantly, we show that CAP2 levels and dimer formation are significantly decreased in transgenic models of Alzheimer disease and in hippocampi of Alzheimer patients, reflecting in aberrant localization of cofilin in spines.

### Impact

This study finally reveals mechanisms controlling activity dependent and structural plasticity through a novel protein partner for cofilin, CAP2. Moreover, the study shows that this pathway is altered in Alzheimer Disease deciphering mechanisms of structural plasticity defective in the disease. Therefore, CAP2 can be considered a marker of the loss of structural plasticity of the synapses, thus paving new ways in the design of synapse-tailored therapeutic strategies and in the identification of new biomarkers of synaptic frailty. This important approach is dramatically needed for Alzheimer Disease therapy and early diagnosis, since it is completely lacking in the drug discovery scenario.

## Expanded View Figure legends

**Figure EV1**

A Representative WB showing the expression of CAP2 in different brain areas (CTX cortex, HIP hippocampus, STR striatum, CB cerebellum, BS brain stem) at postnatal day (P) 0, P7, P14, P18, P21 and P30.

B Representative WB of the levels of markers of the presynaptic (synaptophysin, Syn) and postsynaptic compartment (PSD-95) in total homogenate (HOMO) and postsynaptic fraction (TIF) of human hippocampus. Tubulin is the loading control protein.

**Figure EV2**

A Confocal and STED images of hippocampal neurons fixed and stained for CAP2 (*magenta*) and F-actin (*cyan*). Scale bar = 2 µm.

A. Representative images of WB analysis of the levels of CAP2 and markers of the presynaptic (synaptophysin, SYN) and postsynaptic compartment (PSD-95) in various subcellular compartments (H: Homogenate; S1: low-speed supernatant; P1: nuclei-associated membranes; S2: high-speed supernatant; P2: crude membrane fraction; Syn: synaptosomes; PSD1: Triton Insoluble postsynaptic fraction; PSD2: postsynaptic density fraction).

B. WB images of lysates obtained from COS7 cells transfected with Myc-CAP2 plus scramble sequence (SCR) or four different CAP2-shRNA sequences. Tubulin served as loading reference and RFP antibody as control of transfection. The quantitative analysis confirmed that the sequence D (CAP2-shRNA D) down-regulates CAP2 expression (SCR vs CAP2-shRNA D \**P* = 0.033; 2-tailed unpaired t-test; *n*= 3 independent experiments,)

D QRT-PCR analysis of CAP2 and BDNF in hippocampal neurons infected with either rAAV-CAP2-shRNA or rAAV-SCR (CAP2, SCR vs CAP2-shRNA \*\*\*\**P*< 0.0001, BDNF, SCR vs CAP2-shRNA \*\*\**P*=0007, *n*=10 independent experiments, 2-tailed unpaired t-test)

E Representative WB images of lysates obtained from COS7 cells transfected with CAP2-shRNA plus either Myc-CAP2 or shr-wt-CAP2. RFP antibody served as control for transfection. The blots show that shr-wt-CAP2 is resistant to CAP2-shRNA.

**Figure EV3**

A Domains organization of CAP2 from *M. musculus*.

B Multi-sequence alignment of the CAP1 and CAP2 sequences from *H. sapiens* and *M. musculus* and CAP from *D. discoideumis*. The domains organization is also highlighted in the multi-sequence alignment, following the same color code reported in panel A.

C Representative images of control experiments carried out to verify the specificity of PLA signal revealing EGFP-CAP2/Myc-CAP2 interaction. Neurons were transfected with Myc-CAP2 and EGFP-CAP2 and the PLA assay was carried out in presence of only the primary anti-Myc antibody. In these conditions no PLA signals were generated and detected along MAP2-positive dendrites (*magenta*); scale bar = 5 µm.

**Figure EV4**

A Samples of Streptag-CAP2 and Streptag-(C32G)CAP2 recombinant proteins were loaded onto a non-reducing gel. WB analysis performed with CAP2 antibody showed two bands for the Streptag-CAP2 protein at 84kDa and 168 kDa, corresponding to the monomer and the dimer, while the mutant Streptag-(C32G)CAP2 is detectable as a single band at 84 kDa.

B Representative images of control experiments carried out to verify the specificity of PLA signal revealing Cofilin/Myc-CAP2 interaction. Neurons were transfected with Myc-CAP2 and GFP and the PLA assay was carried out in presence of only the primary antibody for Cofilin (*left panels*) or the anti-Myc antibody (*right panels*). In these conditions no PLA signals were generated and detected along GFP-positive dendrites (*magenta*); scale bar = 5 µm.

C Co-immunoprecipitation assays carried out from homogenate samples of HEK293 cells transfected with Myc-CAP2 or Myc-(C32G)CAP2. Both Myc-CAP2 and Myc-(C32G)CAP2 are able to immunoprecipitate actin.

D Representative WB images of lysates obtained from COS7 cells transfected with CAP2-shRNA plus either Myc-CAP2 or shr-(C32G)-CAP2. RFP antibody served as control for transfection. The blots show that shr-(C32G)-Myc CAP2 is resistant to CAP2-shRNA.

**Figure EV5**

A Samples of rat and human brain Homo and TIF were loaded onto a non-reducing gel. WB analysis performed with CAP2 antibody showed a band corresponding to CAP2 monomer and another band at 106 KDa corresponding to a CAP2 dimer.

B WB representative images of HOMO and TIF samples purified from control and LTD-treated neurons. LTD does not affect CAP2 total levels and localization in the synaptic fraction (CTRL vs LTD *P*>0.05; 2-tailed unpaired t-test, *n*=6 independent experiments)

C WB representative images of TIF samples purified from control and LTD-treated neurons. LTD does not affect CAP2 dimer/monomer ratio in the TIF fraction (CTRL vs LTD, *P*>0.05, 2-tailed unpaired t-test, *n*=6 independent experiments)

D Representative images of PLA showing the proximity between EGFP-CAP2 and either Myc-CAP2 or Myc-(C32G)CAP2 (*white*) along MAP2 positive dendrites (*magenta*) after 30 min of LTD induction; scale bar = 5 µm. LTD induction does not affect CAP2 dimer formation (EGFP-CAP2/Myc-CAP2 vs EGFP-CAP2/Myc-(C32G)CAP2 LTD \*\*\*\**P* < 0.0001, EGFP-CAP2/Myc-CAP2 LTD vs EGFP-CAP2/Myc-(C32G)CAP2 LTD, \*\*\*\**P* < 0.0001, *n*=6-8 neurons from 2 independent experiments, one-way Anova, Bonferroni’s multiple comparisons test).

E Representative images of PLA showing the proximity between CAP2 and either Cofilin (*white*) along MAP2 positive dendrites (*magenta*) after 30 min of cLTP induction; scale bar = 5 µm. LTP induction fosters Cofilin/CAP2 binding (CTRL vs cLTP, \**P* = 0.014, *n*=6-8 neurons from 2 independent experiments, 2-tailed unpaired t-test).

**Figure EV6**

Samples of TIF hippocampi were Ip with a rabbit anti-Cofilin antibody and loaded onto a non-reducing gel. WB analysis revealed the presence of CAP2 monomer and dimer in the immunocomplex. As shown in the right lanes, no signal is detectable when the sample is precipitated without CAP2 antibody. Lanes were run on the same gel but were not contiguous.

## References

Andrianantoandro E & Pollard TD (2006) Mechanism of actin filament turnover by severing and nucleation at different concentrations of ADF/cofilin. Mol. Cell 24: 13–23

Balcer HI, Goodman AL, Rodal AA, Smith E, Kugler J, Heuser JE & Goode BL (2003) Coordinated regulation of actin filament turnover by a high-molecular-weight Srv2/CAP complex, cofilin, profilin, and Aip1. Curr. Biol. 13: 2159–2169

Bamburg JR & Bernstein BW (2016) Actin dynamics and cofilin-actin rods in alzheimer disease. Cytoskeleton (Hoboken) 73: 477–497

Barone E, Mosser S & Fraering PC (2014) Inactivation of brain Cofilin-1 by age, Alzheimer’s disease and γ-secretase. Biochim. Biophys. Acta 1842: 2500–2509

Bernstein BW & Bamburg JR (2010) ADF/cofilin: a functional node in cell biology. Trends Cell Biol. 20: 187–195

Bertling E, Hotulainen P, Mattila PK, Matilainen T, Salminen M & Lappalainen P (2004) Cyclase-associated protein 1 (CAP1) promotes cofilin-induced actin dynamics in mammalian nonmuscle cells. Mol. Biol. Cell 15: 2324–2334

Blalock EM, Geddes JW, Chen KC, Porter NM, Markesbery WR & Landfield PW (2004) Incipient Alzheimer’s disease: microarray correlation analyses reveal major transcriptional and tumor suppressor responses. Proc. Natl. Acad. Sci. U.S.A. 101: 2173–2178

Blanchoin L & Pollard TD (1999) Mechanism of interaction of Acanthamoeba actophorin (ADF/Cofilin) with actin filaments. Journal of Biological Chemistry 274: 15538–15546

Bosch M, Castro J, Saneyoshi T, Matsuno H, Sur M & Hayashi Y (2014) Structural and molecular remodeling of dendritic spine substructures during long-term potentiation. Neuron 82: 444–459

Bourne JN & Harris KM (2008) Balancing Structure and Function at Hippocampal Dendritic Spines. Annu. Rev. Neurosci. 31: 47–67

Braak H & Braak E (1991) Neuropathological stageing of Alzheimer-related changes. Acta Neuropathol. 82: 239–259

Chen LY, Rex CS, Casale MS, Gall CM & Lynch G (2007) Changes in synaptic morphology accompany actin signaling during LTP. J. Neurosci. 27: 5363–5372

Cingolani LA & Goda Y (2008) Differential involvement of beta3 integrin in pre- and postsynaptic forms of adaptation to chronic activity deprivation. Neuron Glia Biol. 4: 179–187

DeKosky ST & Scheff SW (1990) Synapse loss in frontal cortex biopsies in Alzheimer’s disease: correlation with cognitive severity. Ann. Neurol. 27: 457–464

Dinamarca MC, Guzzetti F, Karpova A, Lim D, Mitro N, Musardo S, Mellone M, Marcello E, Stanic J, Samaddar T, Burguière A, Caldarelli A, Genazzani AA, Perroy J, Fagni L, Canonico PL, Kreutz MR, Gardoni F & Di Luca M (2016) Ring finger protein 10 is a novel synaptonuclear messenger encoding activation of NMDA receptors in hippocampus. Elife 5: e12430

Epis R, Marcello E, Gardoni F, Vastagh C, Malinverno M, Balducci C, Colombo A, Borroni B, Vara H, Dell’Agli M, Cattabeni F, Giustetto M, Borsello T, Forloni G, Padovani A & Di Luca M (2010) Blocking ADAM10 synaptic trafficking generates a model of sporadic Alzheimer’s disease. Brain 133: 3323–3335

Field J, Ye DZ, Shinde M, Liu F, Schillinger KJ, Lu M, Wang T, Skettini M, Xiong Y, Brice AK, Chung DC & Patel VV (2015) CAP2 in cardiac conduction, sudden cardiac death and eye development. Sci Rep 5: 17256

Fukazawa Y, Saitoh Y, Ozawa F, Ohta Y, Mizuno K & Inokuchi K (2003) Hippocampal LTP is accompanied by enhanced F-actin content within the dendritic spine that is essential for late LTP maintenance in vivo. Neuron 38: 447–460

Gardoni F, Schrama LH, Kamal A, Gispen WH, Cattabeni F & Di Luca M (2001) Hippocampal synaptic plasticity involves competition between Ca2+/calmodulin-dependent protein kinase II and postsynaptic density 95 for binding to the NR2A subunit of the NMDA receptor. J. Neurosci. 21: 1501–1509

Gu J, Lee CW, Fan Y, Komlos D, Tang X, Sun C, Yu K, Hartzell HC, Chen G, Bamburg JR & Zheng JQ (2010) ADF/cofilin-mediated actin dynamics regulate AMPA receptor trafficking during synaptic plasticity. Nat Neurosci 13: 1208–1215

Henriques AG, Oliveira JM, Carvalho LP & da Cruz E Silva OAB (2015) Aβ Influences Cytoskeletal Signaling Cascades with Consequences to Alzheimer’s Disease. Mol. Neurobiol. 52: 1391–1407

Hild G, Kalmár L, Kardos R, Nyitrai M & Bugyi B (2014) The other side of the coin: functional and structural versatility of ADF/cofilins. European Journal of Cell Biology 93: 238–251

Holtmaat A & Svoboda K (2009) Experience-dependent structural synaptic plasticity in the mammalian brain. Nature Publishing Group 10: 647–658

Horch HW & Katz LC (2002) BDNF release from single cells elicits local dendritic growth in nearby neurons. Nat Neurosci 5: 1177–1184

Hotulainen P & Hoogenraad CC (2010) Actin in dendritic spines: connecting dynamics to function. The Journal of Cell Biology 189: 619–629

Hotulainen P, Llano O, Smirnov S, Tanhuanpää K, Faix J, Rivera C & Lappalainen P (2009) Defining mechanisms of actin polymerization and depolymerization during dendritic spine morphogenesis. The Journal of Cell Biology 185: 323–339

Hubberstey AV & Mottillo EP (2002) Cyclase-associated proteins: CAPacity for linking signal transduction and actin polymerization. FASEB J. 16: 487–499

Jankowsky JL, Fadale DJ, Anderson J, Xu GM, Gonzales V, Jenkins NA, Copeland NG, Lee MK, Younkin LH, Wagner SL, Younkin SG & Borchelt DR (2004) Mutant presenilins specifically elevate the levels of the 42 residue beta-amyloid peptide in vivo: evidence for augmentation of a 42-specific gamma secretase. Hum. Mol. Genet. 13: 159–170

Kasai H, Fukuda M, Watanabe S, Hayashi-Takagi A & Noguchi J (2010) Structural dynamics of dendritic spines in memory and cognition. Trends Neurosci. 33: 121–129

Kim T, Vidal GS, Djurisic M, William CM, Birnbaum ME, Garcia KC, Hyman BT & Shatz CJ (2013) Human LilrB2 is a β-amyloid receptor and its murine homolog PirB regulates synaptic plasticity in an Alzheimer’s model. Science 341: 1399–1404

Korobova F & Svitkina T (2010) Molecular architecture of synaptic actin cytoskeleton in hippocampal neurons reveals a mechanism of dendritic spine morphogenesis. Mol. Biol. Cell 21: 165–176

Koskinen M, Bertling E & Hotulainen P (2012) Methods to measure actin treadmilling rate in dendritic spines. Meth. Enzymol. 505: 47–58

Kumar A, Paeger L, Kosmas K, Kloppenburg P, Noegel AA & Peche VS (2016) Neuronal Actin Dynamics, Spine Density and Neuronal Dendritic Complexity Are Regulated by CAP2. Front Cell Neurosci 10: 180

Lacor PN, Buniel MC, Furlow PW, Clemente AS, Velasco PT, Wood M, Viola KL & Klein WL (2007) Abeta oligomer-induced aberrations in synapse composition, shape, and density provide a molecular basis for loss of connectivity in Alzheimer’s disease. J. Neurosci. 27: 796–807

Liu Y, Xiao W, Shinde M, Field J & Templeton DM (2018) Cadmium favors F-actin depolymerization in rat renal mesangial cells by site-specific, disulfide-based dimerization of the CAP1 protein. Arch. Toxicol. 92: 1049–1064

Marcello E, Epis R, Saraceno C & Di Luca M (2012a) Synaptic dysfunction in Alzheimer’s disease. Adv. Exp. Med. Biol. 970: 573–601

Marcello E, Epis R, Saraceno C, Gardoni F, Borroni B, Cattabeni F, Padovani A & Di Luca M (2012b) SAP97-mediated local trafficking is altered in Alzheimer disease patients’ hippocampus. Neurobiol. Aging 33: 422.e1–10

Marcello E, Gardoni F, Di Luca M & Pérez-Otaño I (2010) An arginine stretch limits ADAM10 exit from the endoplasmic reticulum. J. Biol. Chem. 285: 10376–10384

Marcello E, Gardoni F, Mauceri D, Romorini S, Jeromin A, Epis R, Borroni B, Cattabeni F, Sala C, Padovani A & Di Luca M (2007) Synapse-associated protein-97 mediates alpha-secretase ADAM10 trafficking and promotes its activity. J. Neurosci. 27: 1682–1691

Marcello E, Saraceno C, Musardo S, Vara H, la Fuente de AG, Pelucchi S, Di Marino D, Borroni B, Tramontano A, Pérez-Otaño I, Padovani A, Giustetto M, Gardoni F & Di Luca M (2013) Endocytosis of synaptic ADAM10 in neuronal plasticity and Alzheimer’s disease. J. Clin. Invest. 123: 2523–2538

Matus A (2005) Growth of dendritic spines: a continuing story. Curr. Opin. Neurobiol. 15: 67–72

Mikhaylova M, Bär J, van Bommel B, Schätzle P, Yuanxiang P, Raman R, Hradsky J, Konietzny A, Loktionov EY, Reddy PP, Lopez-Rojas J, Spilker C, Kobler O, Raza SA, Stork O, Hoogenraad CC & Kreutz MR (2018) Caldendrin Directly Couples Postsynaptic Calcium Signals to Actin Remodeling in Dendritic Spines. Neuron 97: 1110–1125.e14

Minamide LS, Striegl AM, Boyle JA, Meberg PJ & Bamburg JR (2000) Neurodegenerative stimuli induce persistent ADF/cofilin-actin rods that disrupt distal neurite function. Nat. Cell Biol. 2: 628–636

Noegel AA, Rivero F, Albrecht R, Janssen KP, Köhler J, Parent CA & Schleicher M (1999) Assessing the role of the ASP56/CAP homologue of Dictyostelium discoideum and the requirements for subcellular localization. J. Cell. Sci. 112 (Pt 19): 3195–3203

Normoyle KPM & Brieher WM (2012) Cyclase-associated protein (CAP) acts directly on F-actin to accelerate cofilin-mediated actin severing across the range of physiological pH. J. Biol. Chem. 287: 35722–35732

Okamoto K-I, Nagai T, Miyawaki A & Hayashi Y (2004) Rapid and persistent modulation of actin dynamics regulates postsynaptic reorganization underlying bidirectional plasticity. Nat Neurosci 7: 1104–1112

Ono S (2013) The role of cyclase-associated protein in regulating actin filament dynamics - more than a monomer-sequestration factor. J. Cell. Sci. 126: 3249–3258

Peche V, Shekar S, Leichter M, Korte H, Schröder R, Schleicher M, Holak TA, Clemen CS, Ramanath-Y B, Pfitzer G, Karakesisoglou I & Noegel AA (2007) CAP2, cyclase-associated protein 2, is a dual compartment protein. Cell. Mol. Life Sci. 64: 2702–2715

Peche VS, Holak TA, Burgute BD, Kosmas K, Kale SP, Wunderlich FT, Elhamine F, Stehle R, Pfitzer G, Nohroudi K, Addicks K, Stöckigt F, Schrickel JW, Gallinger J, Schleicher M & Noegel AA (2013) Ablation of cyclase-associated protein 2 (CAP2) leads to cardiomyopathy. Cell. Mol. Life Sci. 70: 527–543

Penzes P & Vanleeuwen J-E (2011) Impaired regulation of synaptic actin cytoskeleton in Alzheimer’s disease. Brain Res Rev 67: 184–192

Penzes P, Cahill ME, Jones KA, Vanleeuwen J-E & Woolfrey KM (2011) Dendritic spine pathology in neuropsychiatric disorders. Nat Neurosci 14: 285–293

Pontrello CG, Sun M-Y, Lin A, Fiacco TA, DeFea KA & Ethell IM (2012) Cofilin under control of β-arrestin-2 in NMDA-dependent dendritic spine plasticity, long-term depression (LTD), and learning. Proc. Natl. Acad. Sci. U.S.A. 109: E442–51

Quintero-Monzon O, Jonasson EM, Bertling E, Talarico L, Chaudhry F, Sihvo M, Lappalainen P & Goode BL (2009) Reconstitution and dissection of the 600-kDa Srv2/CAP complex: roles for oligomerization and cofilin-actin binding in driving actin turnover. Journal of Biological Chemistry 284: 10923–10934

Racz B & Weinberg RJ (2006) Spatial organization of cofilin in dendritic spines. Neuroscience 138: 447–456

Rush T, Martinez-Hernandez J, Dollmeyer M, Frandemiche ML, Borel E, Boisseau S, Jacquier-Sarlin M & Buisson A (2018) Synaptotoxicity in Alzheimer’s Disease Involved a Dysregulation of Actin Cytoskeleton Dynamics through Cofilin 1 Phosphorylation. J. Neurosci. 38: 10349–10361

Rust MB (2015a) ADF/cofilin: a crucial regulator of synapse physiology and behavior. Cell. Mol. Life Sci. 72: 3521–3529

Rust MB (2015b) Novel functions for ADF/cofilin in excitatory synapses - lessons from gene-targeted mice. Commun Integr Biol 8: e1114194

Rust MB, Gurniak CB, Renner M, Vara H, Morando L, Görlich A, Sassoè-Pognetto M, Banchaabouchi MA, Giustetto M, Triller A, Choquet D & Witke W (2010) Learning, AMPA receptor mobility and synaptic plasticity depend on n-cofilin-mediated actin dynamics. EMBO J. 29: 1889–1902

Sala C & Segal M (2014) Dendritic spines: the locus of structural and functional plasticity. Physiol. Rev. 94: 141–188

Selkoe DJ (2002) Alzheimer’s disease is a synaptic failure. Science 298: 789–791

Shankar GM, Bloodgood BL, Townsend M, Walsh DM, Selkoe DJ & Sabatini BL (2007) Natural oligomers of the Alzheimer amyloid-beta protein induce reversible synapse loss by modulating an NMDA-type glutamate receptor-dependent signaling pathway. J. Neurosci. 27: 2866–2875

Smith KR, Kopeikina KJ, Fawcett-Patel JM, Leaderbrand K, Gao R, Schürmann B, Myczek K, Radulovic J, Swanson GT & Penzes P (2014) Psychiatric risk factor ANK3/ankyrin-G nanodomains regulate the structure and function of glutamatergic synapses. Neuron 84: 399–415

Star EN, Kwiatkowski DJ & Murthy VN (2002) Rapid turnover of actin in dendritic spines and its regulation by activity. Nat Neurosci 5: 239–246

Van Troys M, Huyck L, Leyman S, Dhaese S, Vandekerkhove J & Ampe C (2008) Ins and outs of ADF/cofilin activity and regulation. European Journal of Cell Biology 87: 649–667

Walsh DM, Klyubin I, Fadeeva JV, Cullen WK, Anwyl R, Wolfe MS, Rowan MJ & Selkoe DJ (2002) Naturally secreted oligomers of amyloid beta protein potently inhibit hippocampal long-term potentiation in vivo. Nature 416: 535–539

Yusof AM, Hu N-J, Wlodawer A & Hofmann A (2005) Structural evidence for variable oligomerization of the N-terminal domain of cyclase-associated protein (CAP). Proteins 58: 255–262

Yuste R & Bonhoeffer T (2001) Morphological changes in dendritic spines associated with long-term synaptic plasticity. Annu. Rev. Neurosci. 24: 1071–1089

Zhou Q, Homma KJ & Poo M-M (2004) Shrinkage of dendritic spines associated with long-term depression of hippocampal synapses. Neuron 44: 749–757

